# Discovery of a FANCD2-interacting protein motif (DIP-box) linking DNA Damage Response processes

**DOI:** 10.64898/2026.02.24.707417

**Authors:** Zhenbo Cao, Giuseppina R Briola, Clara Ionita, James Streetley, Helen Walden, Martin Luke Rennie

**Affiliations:** School of Molecular Biosciences, College of Medical Veterinary and Life Sciences, University of Glasgow; Glasgow, UK; Scottish Centre for Macromolecular Imaging, University of Glasgow, Glasgow, UK

## Abstract

Repair of DNA damage is critical to genome integrity and the DNA clamp, FANCD2, is a major player in the response to interstrand crosslinks tying the two DNA strands together. It has long been suggested FANCD2 recruits other factors to coordinate repair these crosslinks, however it remains unclear if FANCD2 directly mediates this and how this is achieved. Here we show that FANCD2 has a conserved acidic region that directly recognises established DNA repair proteins containing a short linear motif. We discover new candidate interactors engaging with FANCD2 in a similar mode that are involved in histone modification, and RNA processing. Our data demonstrate FANCD2 is a hub for protein-protein interactions providing a structural explanation for FANCD2’s central role in repair of DNA interstrand crosslinks.

**Graphical Abstract:** 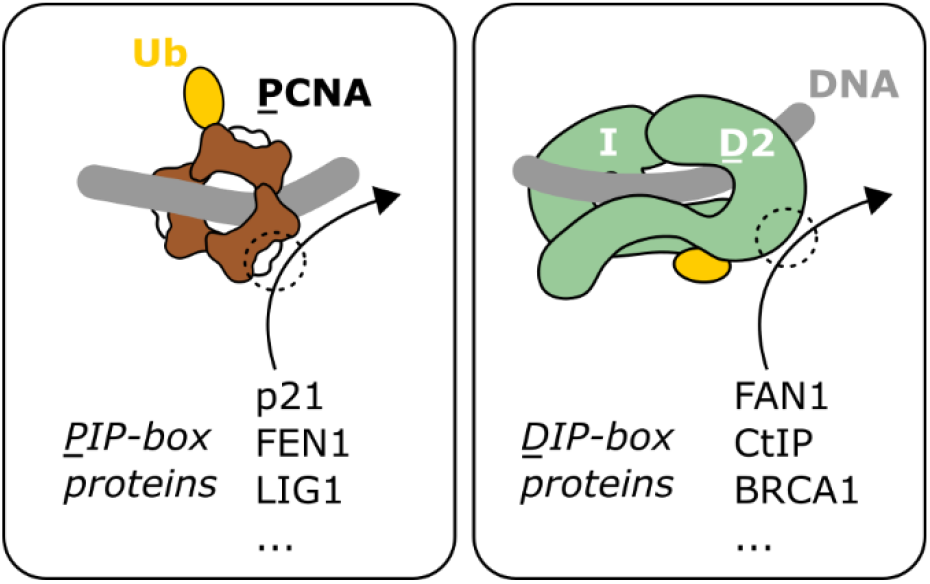

## Introduction

DNA damage response (DDR) pathways facilitate repair of DNA lesions that arise from both endogenous and exogenous sources (*1*). These pathways involve overlapping sets of proteins and have extensive redundancy that allows robust repair of damage. The Fanconi Anemia pathway is one such pathway and helps to repair interstrand crosslinks (ICLs) by utilizing homologous recombination (HR) processes (*2, 3*). ICL repair is a complex process involving nuclease incisions that unhook the crosslink lesion but generate DNA double-strand breaks (DSBs), the ends of which are resected to allow HR (*3*). These processes are carefully orchestrated by numerous DDR proteins and their post-translation modifications (PTMs) (*1*–*3*). A key step in the Fanconi Anemia pathway is the specific mono-ubiquitination of the Fanconi Anemia complementation group D2 (FANCD2) protein. This PTM leads FANCD2 to colocalize with other DDR proteins in nuclear foci (*4*–*12*), however the structural underpinnings of the interactions within these foci are unclear.

It was initially hypothesised that the ubiquitin conjugated to FANCD2 could act to physically recruit DDR proteins that have ubiquitin-interacting domains (*13*), however subsequent structural and biochemical studies suggest this is unlikely to be the case. FANCD2 forms a hetero-dimer with its paralog, FANCI, and structures of the FANCI-FANCD2 complex show that it is a DNA clamp, with mono-ubiquitination acting to stabilize the complex on DNA (*14*–*18*). In these structures FANCI sequesters a substantial portion of the ubiquitin surface (*15, 16*) and FANCD2’s ubiquitin has reduced accessibility for some deubiquitinases when in the context of the FANCI-FANCD2 complex (*17*). Furthermore, it has been observed that the interaction between the FANCI-FANCD2 complex and several candidate interactors is not drastically altered by mono-ubiquitination (*18*). These studies suggest mono-ubiquitination traps the FANCI-FANCD2 complex on the DNA at sites of damage rather than directly recruiting other DDR proteins.

Several DDR proteins have been shown to co-localize with FANCD2 (*4*–*12*). CtIP (C-terminal binding protein 1-interacting protein) is one such protein (*10*–*12*), and activates the MRE11-RAD50 nuclease for end resection at DSBs. CtIP forms an elongated tetramer and can recognise DNA fork structures (*19*), and is directed to DNA damage foci by mono-ubiquitinated FANCD2 (*10*–*12*). FAN1 (Fanconi-associated nuclease 1) is another proposed interactor of the FANCI-FANCD2 complex, and is a nuclease that can cleave branched DNA structures that arise during ICL excision (*6*–*9*) and prevent chromosome abnormalities at stalled replication forks (*20*). Like CtIP, FAN1 is directed to DNA damage foci by mono-ubiquitinated FANCD2, dependant up a Ubiquitin-Binding Zinc finger (UBZ) domain of FAN1. BRCA1 (Breast cancer type 1 susceptibility protein) is also reported to co-localize with FANCD2 in nuclear foci (*4, 5*). BRCA1 ubiquitinates histones and CtIP to facilitate end resection (*21*). More broadly, FANCD2 has been suggested to have a role in histone (*22*) and R-loop processing (*23, 24*). Finally, to remove the ubiquitin signal and presumably unlock FANCD2 from the DNA for recycling of the clamp, multi-faceted interactions with the deubiquitinase, USP1, are required (*25*– In particular, a short region of the disordered N-terminal extension of USP1 is critical to recognise FANCD2 (*26, 27*). How FANCD2 physically connects to these various proteins and pathways is not well understood.

Proliferating Cell Nuclear Antigen (PCNA) shares similarities with FANCD2 in that it is a DNA clamp involved in the DDR, is specifically mono-ubiquitinated, and is deubiquitinated by USP1 (*28*–*30*). PCNA is a central player in DNA replication and repair having a diverse range of functions, in particular as a processivity factor for polymerases (*28*). Mono-ubiquitination of PCNA recruits specialized polymerases to bypass DNA damage (*29*). PCNA is also well-established to interact with numerous partner proteins through a PCNA-interacting motif (PIP-box), facilitating its diverse functions (*28*). These are generally short linear motifs (SLiMs) within disordered regions of the interactor. Here we show that various interactors of FANCD2 are mediated via SLiMs, in a similar fashion to PCNA, explaining the central role of FANCD2 in ICL repair.

Using a combination of cryoEM, AlphaFold, and biochemical reconstitution, we identify a FANCD2 interacting motif in several DDR proteins, which we refer to as a D2-interacting protein box (DIP-box), and the corresponding site on FANCD2 to which DIP-boxes bind. We show that CtIP, FAN1, and USP1 all contain a DIP-box consisting of a leucine surrounded by positively charged residues that interact with a conserved acidic region on FANCD2. We further predict DIP-boxes in nine other proteins, including BRCA1 and those involved in histone modifications and RNA processing. This work provides a structural basis for the various cellular functionalities of FANCD2.

## Results

### Structural basis of FAN1 and CtIP interaction with FANCD2 resolved by cryoEM

We sought to establish a reconstituted setup to probe the structural basis of FANCD2-CtIP and FANCD2-FAN1 interactions. As a first step we generated AlphaFold3 (*31*) predictions of FANCD2 with FAN1 and with CtIP, which highlight a short section of each protein with predicted aligned error (PAE) scores much lower (more confident) than the surrounding residues, indicating a potential interaction (Figure 1A, S1A). For CtIP, this region is consistent with previous data in the presence of cellular components (*10, 12*). It has been shown that for predictions involving SLiMs, the use of fragments of protein sequences can increase sensitivity (*32*). As such we generated further AlphaFold3 predictions using fragments of FAN1 or CtIP around the candidate site (Figure S1B). This approach improved the confidence scores and suggested the UBZ-domain of FAN1 (residues 41-69) may interact directly with FANCD2.

**Figure 1:**
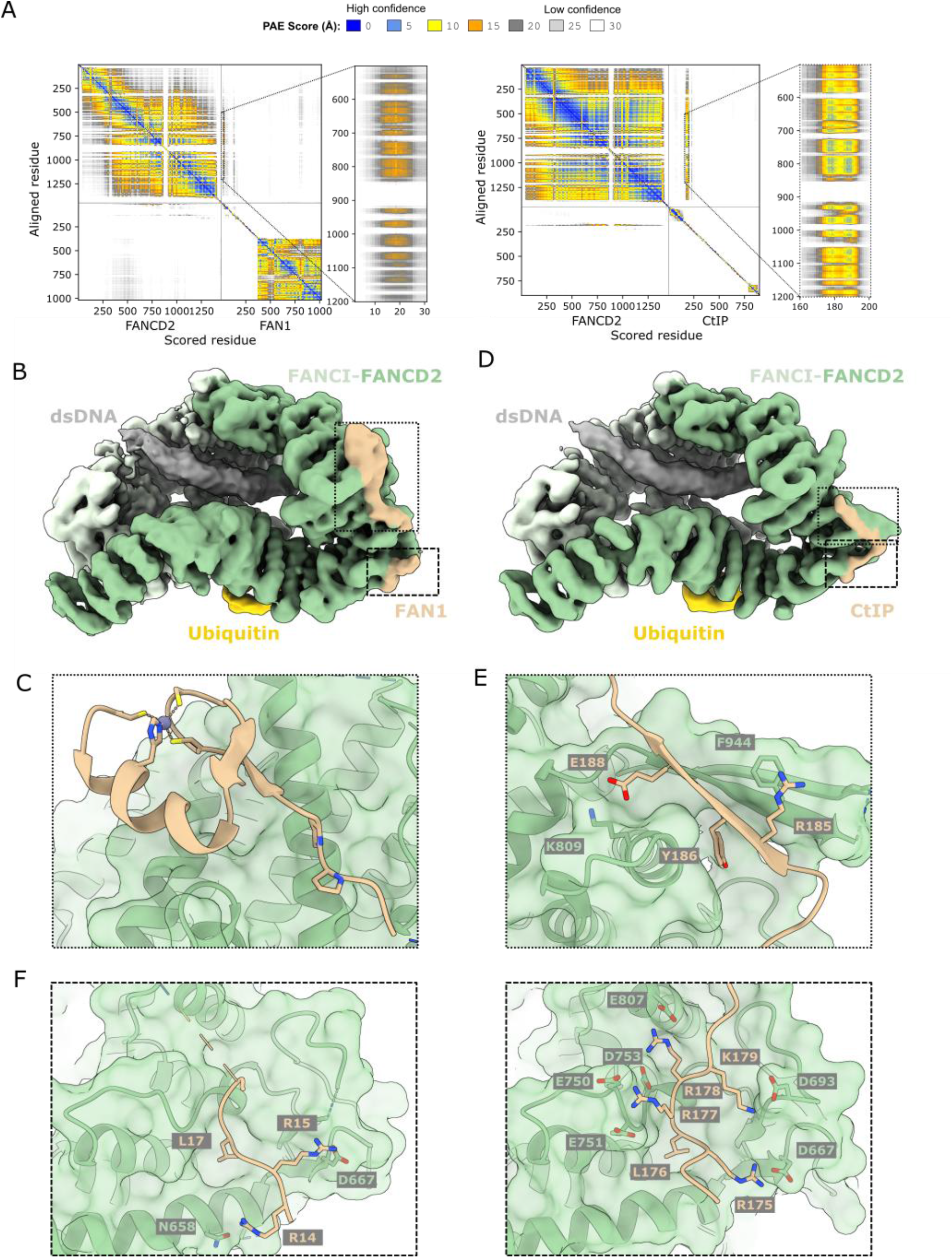
FAN1 and CtIP interact with FANCD2 in a partially overlapping bipartite manor. (A) AlphaFold3 PAE plot of FANCD2 with FAN1 (left) and CtIP (right). Insets highlight the small regions of confident interaction. (B) CryoEM consensus reconstruction of the FANCI-FANCD2^Ub^-FAN1^1-71^ complex. (C) Refined model of the UBZ interaction with FANCD2. (D) CryoEM consensus reconstruction of the FANCI-FANCD2^Ub^-CtIP^173-193^ complex. (E) Refined model of the β-sheet interaction with FANCD2. (F) Refined model of FAN1 and FANCD2 (left), and CtIP and FANCD2 (right) at the acidic site.

To understand how FAN1 recognises the FANCI-FANCD2 clamp, we designed a truncation of FAN1 encompassing the UBZ and other candidate interaction site (FAN1^1-71^) and reconstituted it with FANCI and mono-ubiquitinated FANCD2 and double-stranded DNA (dsDNA) for cryoEM analysis. We were able to reconstruct the FANCI-FANCD2^Ub^-dsDNA assembly containing the FAN1 fragment to ∼3.5 Å (Figure S2, Table S1). The overall complex resembles structures of the closed conformation of mono-ubiquitinated FANCI-FANCD2, with the ubiquitin sequestered by FANCI (*15*–*17, 33, 34*) (Figure 1B). We identified density corresponding to FAN1 in the helical domain (HD) and C-terminal helical repeat domain (CTD) of FANCD2. We refined an AlphaFold3 model of FANCD2-FAN1 into a cryoEM map focused on this region to generate an atomic model of the interaction (Figure S2). The UBZ domain of FAN1 and a pair of preceding prolines (residues 38-64) interacts with the CTD of FANCD2 (Figure 1C).

To understand how CtIP recognises the FANCI-FANCD2 clamp, we designed a peptide fragment of CtIP encompassing the candidate FANCD2 interaction site (CtIP^173-193^) and reconstituted it with FANCI and mono-ubiquitinated FANCD2 and double-stranded DNA (dsDNA) for cryoEM analysis. We were able to reconstruct the FANCI-FANCD2^Ub^-dsDNA assembly containing the CtIP fragment to approximately 4 Å (Figure S3, Table S1). The overall complex again resembles structures of the closed conformation of mono-ubiquitinated FANCI-FANCD2, with the ubiquitin sequestered by FANCI (*15*– *17, 33, 34*) (Figure 1D). We identified density corresponding to CtIP in the HD of FANCD2. We refined an AlphaFold3 model of the FANCD2-CtIP interaction into a cryoEM map focused on this region to generate an atomic model of the interaction site (Figure S3). A short stretch of residues of CtIP, 184-189, form an anti-parallel β-sheet with FANCD2 (residues 940-958) (Figure 1E). On this strand a tyrosine (Y186) also contributes hydrophobic interactions and just downstream a glutamate of CtIP (E188) interacts with a lysine of FANCD2. This β-sheet structure of FANCD2 is not observed in any other structures of FANCD2 and driven by a substantial conformational change (Figure S4) (*15, 25*). To form the β-sheet, residues 938-944, forming part of the FANCD2 core in other structures, extend out and residues 945-958 become ordered. This rearrangement suggests CtIP may act to stabilize this region or provide a platform for additional protein-protein interactions.

Both FAN1 and CtIP have an additional interaction point with FANCD2, upstream in sequence from the UBZ and β-strand, respectively (Figure 1F). While the UBZ and β-sheet interaction sites are distinct, the upstream interaction point is strikingly similar. Both interactions involve a leucine, L17 in FAN1 and L176 in CtIP, buried into a hydrophobic pocket of FANCD2. We refer to this as the leucine anchor. An arginine nearby, R15 in FAN1 and R175 in CtIP, interacts with D667 of FANCD2. For CtIP, additional positively charged residues on the other side of the leucine anchor contact negatively charged residues on FANCD2. In FAN1, there are additional positively charged residues on the other side of the leucine anchor that may contribute to the interaction; however they are not clearly resolved in our cryoEM reconstruction. These data reveal bipartite interaction modes with FANCD2, with partial overlap of the interaction sites at an acidic site on FANCD2.

### SLiM interactions with FANCD2 underpin its role in ICL repair

The acidic interaction site on FANCD2 that both FAN1 and CtIP bind (Figure 1F) is also implicated in USP1 interaction with FANCD2 (*27*). In the case of USP1, the interacting region lies in the disordered N-terminal extension which is important for efficient FANCD2 deubiquitination (*26*). In previous AlphaFold predictions of FANCD2 with USP1, the leucine anchor of USP1, L23, that docks with FANCD2 is preceded by an arginine, R22, that interacts with D667 of FANCD2 (*27*), similar to that observed in the FAN1 and CtIP experimental structures. Both R22 and L23 are necessary for efficient deubiquitination of FANCD2 by USP1 and there is a lysine, K26, just downstream that contributes to a lesser extent (*26*). USP1 appears to lack the CTD and β-sheet interaction points with FANCD2. However, USP1 forms a complex with an activating partner protein, UAF1, which in turn interacts with FANCI (*25*). Therefore, although USP1 lacks the additional interaction points with FANCD2 identified here, its interaction is supplemented by the partner proteins UAF1 and FANCI.

In the absence of FANCD2 the leucine anchor and surrounding positively charged residues of FAN1, CtIP, and USP1 are predicted with low plDDT scores (AlphaFold Database) and so likely to be unstructured (*35, 36*). Therefore, this interaction point is consistent with a SLiM interaction mode. SLiMs that interact with an overlapping site of the PCNA DNA clamp are referred to as PIP-boxes, so we refer to the leucine anchor flanked by positively charged residues within FANCD2 interactors as a DIP-box (D2-interacting protein box).

We examined the importance of charge of the acidic site on FANCD2 for USP1’s DIP-box interaction using a deubiquitination reporter assay. This assay is based on the requirement of this interaction within the N-terminal extension of USP1 that contains the DIP-box for efficient deubiquitination of FANCD2 (*26*). We mutated acidic residues on FANCD2, focusing on residues either side of the leucine anchor, L23: D667 on FANCD2 contacting R22 on USP1, and E750 and E751 predicted to be adjacent to K26. We compared the activity of USP1-UAF1 on wild-type FANCD2 versus D667R, E750A/E751K, and D667R/E750A/E751K FANCD2 mutants. We first specifically mono-ubiquitinated wild-type

FANCD2 and the mutants at K561 using our established reconstituted system (*37, 38*). The FANCD2 mutants were nearly completely mono-ubiquitinated, suggesting the mutants were folded similarly to the wild-type protein (Figure 2A). We then added USP1-UAF1 and monitored deubiquitination over time. The D667R and E750A/E751K mutants resulted in a reduction in deubiquitination rates, while the combined D667R/E750A/E751K mutant was reduced further. The additive effect of these mutations supports USP1 DIP-box binding to FANCD2 through multivalent interactions flanking the leucine anchor.

**Figure 2:**
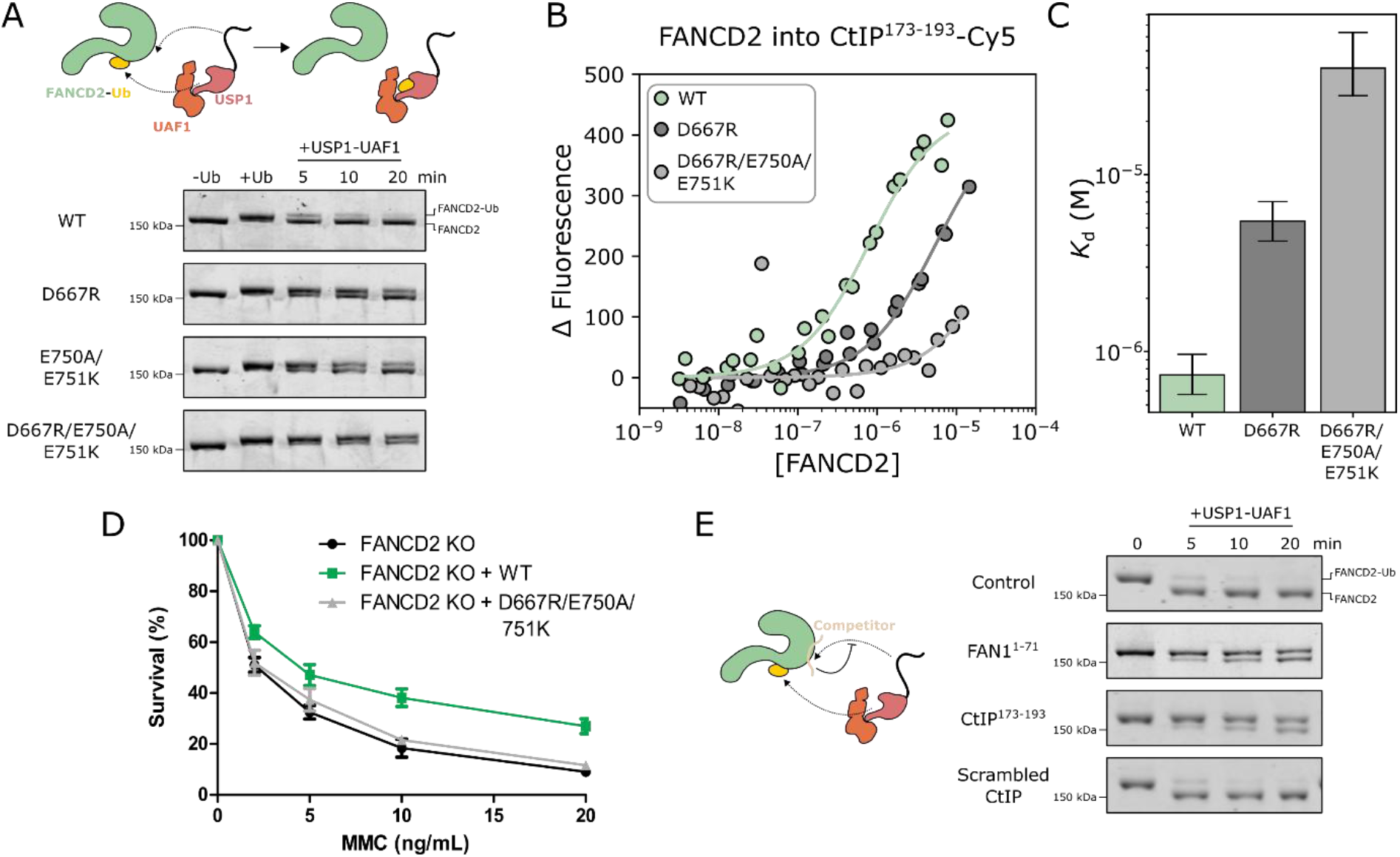
Interaction between the acidic site and DIP-box is important for FANCD2 recognition. (A) Deubiquitination assays for acidic site mutants. “-Ub” refers to purified FANCD2 before the ubiquitination machinery is added, “+Ub” refers to the sample after incubation with the ubiquitination machinery yielding FANCD2 with K561 linked mono-ubiquitin. At least two technical replicates were performed. (B) Fluorescence measurements of wild-type FANCD2 or acidic site mutants titrated into CtIP peptide. Two replicates were performed. (C) Fitted binding affinities. (D) Survival assays for acidic site mutants. U2OS FANCD2 knockout cells transfected with wild-type or mutant FANCD2-GFP were treated with the indicated doses of MMC for 16 hours, and the survival was determined after 7 days. Data are presented as the mean ± SD (n = 5). (E) Competition deubiquitination assays with purified wild-type FANCD2 with K561 linked mono-ubiquitin. USP1-UAF1 at 100 nM was used for all deubiquitination experiments with mono-ubiquitinated FANCD2 substrate at 1 μM. Competitors were included at 4 μM where indicated. At least two technical replicates were performed.

To quantify the FANCD2-CtIP binding and verify the importance of charge for this interaction, we deployed a fluorescence assay (Figure 2B). We used the peptide fragment of CtIP encompassing the FANCD2 binding region (CtIP^173-193^), with a Cy5 label at the C-terminal lysine. Addition of FANCD2 resulted an increase in fluorescence, consistent with binding triggering protein-induced fluorescence enhancement (*17, 39*). Titration of FANCD2 suggests a binding affinity slightly tighter than 1 μM (Figure 2C). We also measured binding to the aforementioned mutants. The single charge reversal, D667R reduced the affinity approximately 10-fold, while the triple mutation, D667R/E750A/E751K, reduced the affinity approximately 50-fold. This additive effect is consistent with multivalent interactions either side of the leucine anchor facilitating the interaction with FANCD2, similar to the deubiquitination data for USP1.

We next determined the biological effect of disrupting DIP-box interaction in U2OS cells. With the FANCD2 triple mutant D667R/E750A/E751K clearly disrupting DIP-box interactions but retaining the ability to be mono-ubiquitinated *in vitro*, we transfected U2OS FANCD2 KO cells to express this mutant or wild-type FANCD2 and measured their survival when treated with the DNA crosslinker, mitomycin C (MMC) (Figure 2D). Transfection of wild-type FANCD2 resulted in greater resistance to MMC compared to non-transfected FANCD2 KO cells, whereas transfection with the D667R/E750A/E751K mutant did not improve resistance to MMC. These data suggest that the DIP-box binding site is crucial for FANCD2’s role in repair of ICLs.

### FAN1 and CtIP compete with USP1 for binding to FANCD2 in vitro

Given that FAN1, CtIP, and USP1 can bind the acidic site of FANCD2 we hypothesised that FAN1 and CtIP may impair USP1-mediated deubiquitination of FANCD2. To test this hypothesis, we repeated the deubiquitination reporter assay but this time including FAN1^1-71^ or CtIP^173-193^ as competitors for USP1 binding to FANCD2. At 4-fold excess of FANCD2^Ub^, FAN1 or CtIP slowed the deubiquitination reaction (Figure 2E). In contrast, scrambled CtIP peptides showed negligible inhibition, even at 50-fold excess of FANCD2, suggesting charge alone does not drive the interaction (Figure 2E, Figure S5). Overall, these data are consistent with our structural models and reveals a mutually exclusive interaction point between USP1 and FAN1 or CtIP for binding to the same site on FANCD2. This competition may act to mitigate premature deubiquitination of FANCD2 while it is still functioning with another interactor.

### The DIP-box binding site on FANCD2 is a conserved hub for protein-protein interactions

The well-established functional sites of FANCD2, for mono-ubiquitination and FANCI interaction, are highly conserved among various eukaryotes (Figure 3A). The DIP-box binding site is also well-conserved (Figure 3A), supporting its functional importance. This site is within an extensive patch of negative density on FANCD2 (Figure 3B). We hypothesized that other proteins involved in DNA repair may also utilize this region to recognise FANCD2.

**Figure 3:**
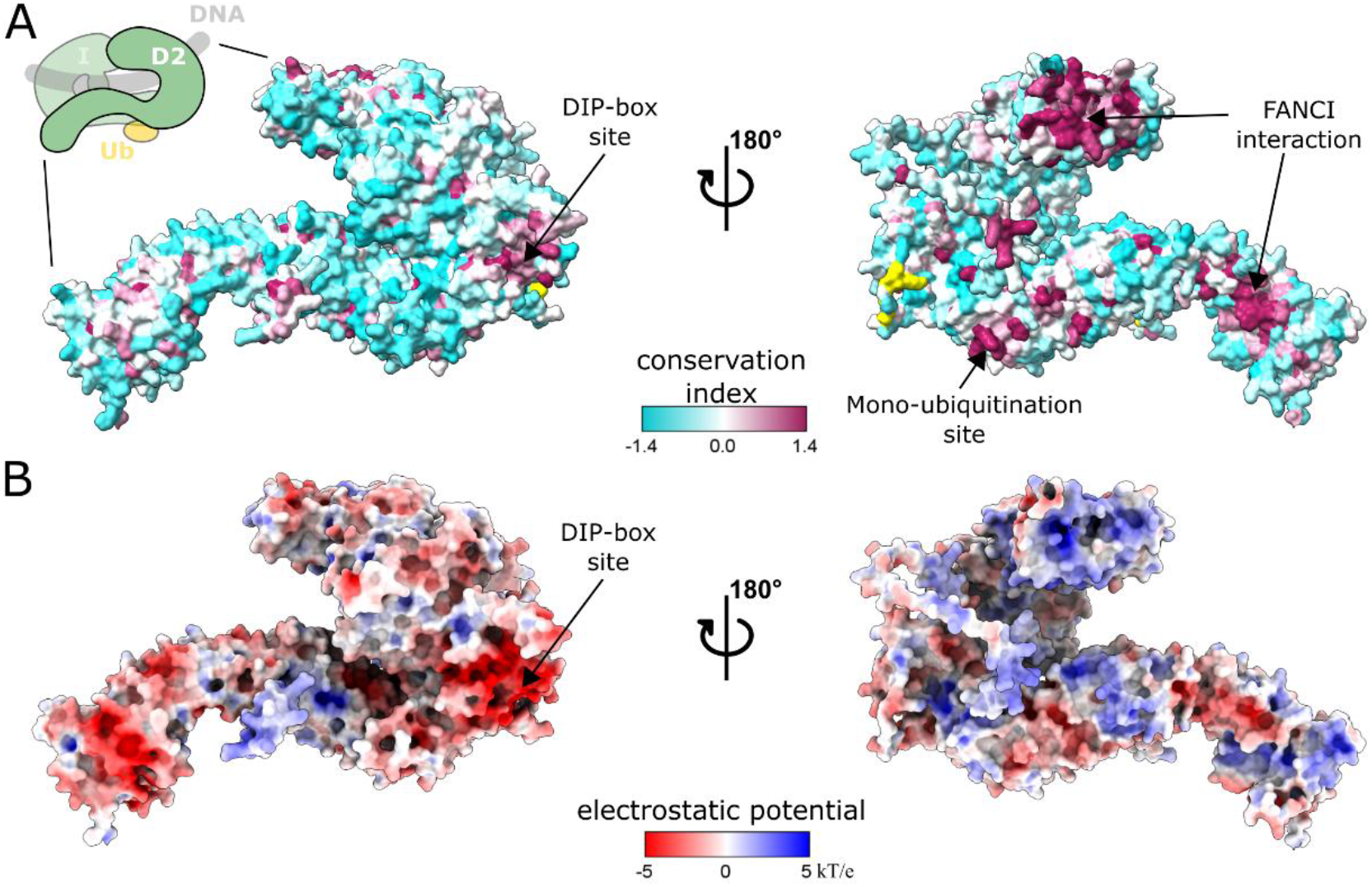
A conserved acidic region of FANCD2 mediates FAN1, CtIP, and USP1 interactions. (A) FANCD2 coloured by sequence conservation index (more positive implying greater conservation). Conservation was quantified by the AL2CO entropy-based measure (*40*) applied to a multiple sequence alignment of Quest for Orthologues species obtained from ProViz (*41*). Functional regions are highlighted. Residues with no assigned conservation index are coloured yellow. (B) FANCD2 surface coloured by electrostatic potential using the APBS web server (*42*). For clarity, the disordered N- and C-termini are not shown.

We sought to identify new candidate DIP-boxes by combining sequence information from the DIP-boxes we have identified, functional information from UniProt (*43*), and co-evolutionary information via AlphaFold3 (*31*). We focused on the leucine anchor that is conserved for the three identified interactors. Given the small number of sequences, we were unable to reliably define a strict motif. Our assays suggest that basic residues that interact with acidic residues on FANCD2 may be required within a short window on both sides of the leucine anchor (Figure 2A-C). We further assumed that leucine anchors should be within unstructured regions in the absence of FANCD2. However, these criteria alone were insufficient to reduce the candidates to a number we considered tractable for further analysis (<100). We therefore explored various criteria on a seven-residue window either side of the leucine anchor and searched human proteins on UniProtKB with experimental or predicted evidence of nuclear localization meeting these criteria. Criteria included a minimal number of positively charged residues, restricting the number of hydrophobic and acidic residues, and restricting to proteins reported to be involved in DNA repair or homologous recombination. Further details are provided in the methods. Our refined criteria yielded 47 candidate sequences from 39 different proteins. We then used AlphaFold3 to predict complexes with FANCD2 of the regions identified with approximately 50-100 residues flanking the region (Table S2). We assumed the confidence metrics for the pooled predictions contained two clusters – one for those with confident predictions and one for those with non-confident predictions. We used K-means clustering on the minimum PAE (minPAE) score and the interface predicted Template Modelling (ipTM) score, respectively, with k=2 for each metric, to assign each prediction to one of these clusters (Figure 4A). We considered those predictions clustering with low minPAE scores and high ipTM scores as confident predictions. All three validated FANCD2 interactors were in this confident cluster, supporting the approach, as well as nine new candidates (Figure 4B, Figure S6). None of the nine new candidates contain confidently predicted supplementary interaction sites like those observed for the CtIP β-strand or the FAN1 UBZ (Figure 4B, Figure S6).

**Figure 4:**
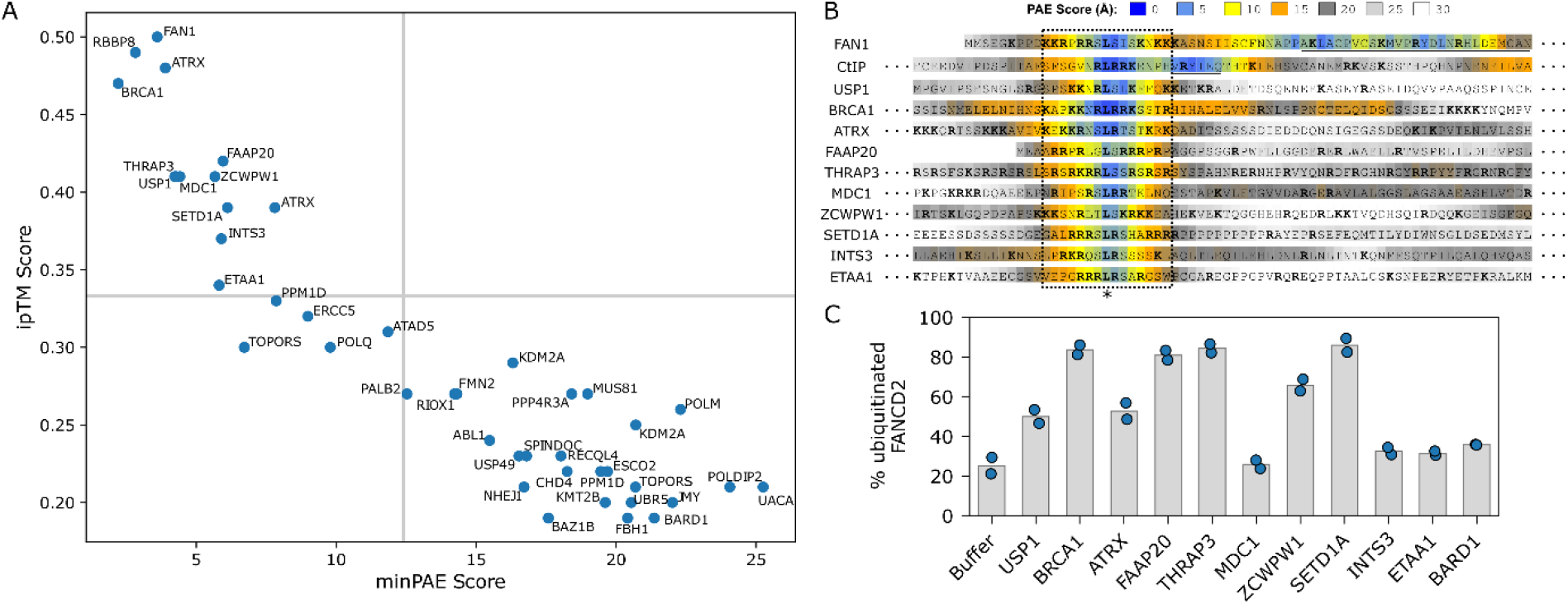
Identification of candidate DIP-boxes by motif search and AlphaFold predictions. (A) Clustering analysis of candidate FANCD2 interactors from motif searching. Thresholds determined by K-means (k=2) for ipTM and minPAE scores respectively are shown as grey lines. Predictions were considered confident if both thresholds were passed (top left quadrant). Some proteins are duplicated due to the presence of multiple candidate motifs within the sequence. (B) PAE confidence scores for each interactor fragment with respect of FANCD2. Each interactor residue is coloured by the lowest numerical PAE value across all FANCD2 residues (0 Å indicating a very confidently predicted position with respect to at least one residue in FANCD2, 30 Å indicating a poorly predicted position). The FAN1 UBZ and CtIP β-sheet interaction points are underlined. Sequences are aligned by the leucine anchor (*). Arginine and lysine residues are highlighted in bold text. (C) Deubiquitination assays with peptides of DIP-box candidates as included as competitors. USP1-UAF1 enzyme at 100 nM was used for all experiments with mono-ubiquitinated FANCD2 substrate at 1 μM and competitors, where included, at 4 μM. Two technical replicates were performed.

To experimentally validate these interactors, we deployed our deubiquitination reporter assay, including peptides for each candidate DIP-box as competitors (Figure 4C). BRCA1, FAAP20, THRAP3, and SETD1A all dramatically reduced the deubiquitination levels consistent with blocking USP1 access to the DIP-box binding site. ATRX and ZCWPW1 had moderate levels of inhibition similar to a USP1 DIP-box peptide. MDC1, INTS3, and ETAA1 had very low levels of inhibition, similar to BARD1, which was in the low confidence cluster. Of the candidates that had a strong effect on deubiquitination, BRCA1 and FAAP20 have established roles in the Fanconi Anemia pathway (*4, 44*), THRAP3 is involved in R-loop processing (*45*), and SETD1A in histone processing (*46*). The moderate inhibitors, ATRX and ZCWPW1, are both involved in histone processes. The particular acidic residues on FANCD2 and total number of electrostatic interactions vary between these predicted structures (Figure S7). While these interactions require further follow-up, it appears the DIP-box binding site unifies FANCD2’s various functions in genome maintenance.

## Discussion

FANCD2 has long been established to co-localize with FAN1 (*6*–*8*) and CtIP (*10*–*12*) in nuclear foci, however the structural details of such interactions, including whether they are mediated by direct interactions have remained elusive. Here we have demonstrated a direct physical interaction between FANCD2 and each of these two proteins through a newly identified DIP-box. We show that disruption of the DIP-box binding site on FANCD2 negates its protective role to DNA crosslinkers. We have also identified BRCA1, FAAP20, THRAP3, and SETD1A as likely DIP-box containing proteins, interacting with FANCD2 in a similar manner. Of these four interactors only BRCA1 and FAAP20 are identified as a potential interactors of FANCD2 in an AlphaFold screen of genome maintenance proteins and even then these have a very low Structure Prediction and Omics informed Classifier (SPOC) scores (≤0.333), indicating a high false discovery rate (*47*). This may reflect that the DIP-boxes are within disordered regions with small interaction regions that are difficult to identify in screens of full-length proteins. Overall, the various interactors we have identified suggest FANCD2 is a hub, similar to PCNA, recruiting other factors to sites of DNA damage and repair.

BRCA1 is well established in homologous recombination and the interaction with FANCD2 provides a direct physical link between the Fanconi Anemia pathway and homologous recombination. It further provides a structural basis for BRCA1 and FANCD2 colocalization in DNA repair foci (*4*). FAAP20 is part of the Fanconi Anemia core complex (FA-CC) which mono-ubiquitinates FANCI and FANCD2 (*44*). The interaction between FAAP20 and FANCD2 may assist in the recruitment of the FANCI-FANCD2 DNA clamp to the FA-CC. FANCD2 is proposed to interact with SETD1A, an enzyme that methylates histones to protect the replication fork during DNA damage (*46*). Our discovery of a direct interaction between these two proteins supports this association and provides a structural basis for the interaction. Finally, FANCD2 has been reported to recruit RNA processing factors to R-loops (*48*). THRAP3 is involved in mRNA splicing and promotes R-loop resolution (*45*), however was not previously reported to be recruited by FANCD2. The direct interaction between FANCD2 and THRAP3 that we have discovered provides a physical basis for the recruitment of THRAP3 to R-loops.

FAN1 and CtIP are both recruited to DNA repair foci by mono-ubiquitinated FANCD2 and involved in cutting specific DNA structures, suggesting FANCD2 plays a critical role in coordinating nuclease activity. Our data reveals the structural mechanisms through which they are recruited. FAN1’s UBZ has been shown to be important for the interaction with FANCD2. It was initially thought that ubiquitin mediated the interaction, however our data suggests that the UBZ directly contacts FANCD2 itself, positioned on the opposite side of FANCD2 to the conjugated ubiquitin (Figure 1B). CtIP is well-established to bind to MRE11-RAD50 which possesses end resection nuclease activity. This interaction site of CtIP (residues 792-869 (*49*)) is distinct from the DIP-box and the β-strand sites (residues 173-193) which we show interact with FANCD2. CtIP has also been shown to tightly interact with forked DNA structures (*19*), while the FANCI-FANCD2 complex appears to stall at single-strand/double-strand DNA junctions (*34*). These suggest the possibility a super-assembly between FANCI, FANCD2, CtIP, MRE11, and RAD50 may form on the DNA during end resection.

Despite the large available surface area of FANCD2 and FANCI available for interactions, all of these proteins bind the same site of FANCD2, a conserved acidic region, via a SLiM that we refer to as a DIP-box. The DIP-box consists of a leucine surrounded by basic residues. Although we were unable to define a simple SLiM motif, scrambling of the DIP-box sequence of CtIP, disrupted the interaction with FANCD2. This suggests charge alone is not the primary driver of the interaction and that the sequence order is important. Further, the DIP-box sequences we identified are very similar to nuclear localization sequences (NLSs). Indeed, for BRCA1 the DIP-box has been identified as one of multiple NLS signatures, however it does not appear to be the primary one (*50*). Furthermore, there is a variant with conflicting classifications of pathogenicity in this region of BRCA1 (NM_007294.4(BRCA1):c.1819A>G (p.Lys607Glu)

https://www.ncbi.nlm.nih.gov/clinvar/variation/224430/). Our models suggest this mutation could disrupt the interaction with FANCD2. We speculate that the DIP-box sequences may have diverged from NLS sequences to gain function to interact with FANCD2.

The DIP-box for both FAN1 and CtIP is supplemented by additional interaction modules that bind outside of the negative region on FANCD2 (Figure 1). This is also the case for USP1 to some extent, as the USP1-UAF1 enzyme complex has an extensive interface between UAF1 and FANCI (*25*). It is possible that the other FANCD2 interactors bind to other regions of FANCD2 via interactions outside of the fragments we have used or partner proteins that supplement their DIP-box interaction.

The various interactors of the DIP-box binding site of FANCD2 raises the question of whether these are involved in a sequential or competitive mechanism in the cellular environment (Figure 5). A sequential mechanism could involve a cooperative hand-off from one binding partner to the next to facilitate efficient DNA repair. A competitive mechanism, on the other hand, could involve the different interactors competing for FANCD2 binding to activate different sub-pathways in DNA repair. Given that FANCD2 forms foci during repair it may also be that multiple interactors are present simultaneously in a concerted mechanism (Figure 5).

**Figure 5:**
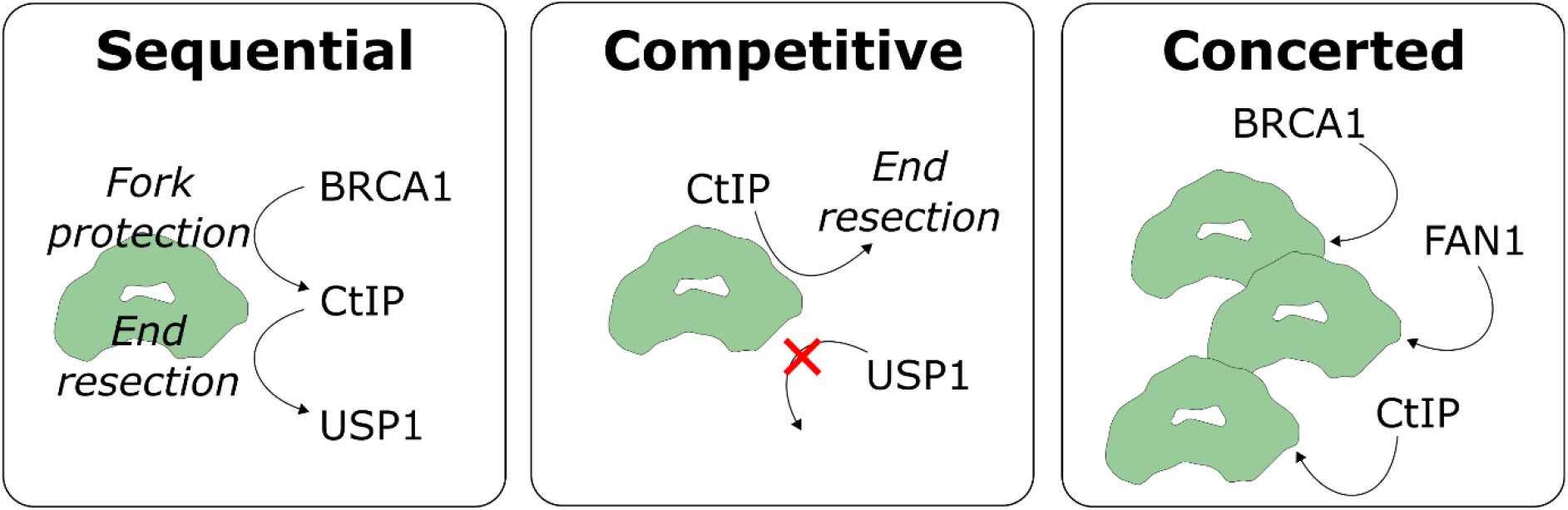
Alternative mechanisms of FANCD2 cooperation with DNA repair machinery. The FANCI-FANCD2 heterodimer is represented as a green blob.

It has been shown that both FANCI and FANCD2 are required for efficient FAN1 and CtIP recruitment to DNA damage sites (*6*–*12, 51*), while BRCA1 is required for efficient FANCD2 recruitment (*4, 11, 52*). This is consistent with a sequential mechanism in which BRCA1 first recruits FANCD2 to DNA damage foci, then FANCD2 recruits FAN1 and/or CtIP for nucleolytic activity, and finally USP1 removes the mono-ubiquitin signal (Figure 5). Interestingly, the supplementary interaction points for FAN1, CtIP, and USP1 are all distinct from one-another, which could facilitate a sequential mechanism involving a cooperative hand-off of FANCD2 to the next interactor. This could be achieved by BRCA1-bound FANCD2 recruiting CtIP via the β-sheet interaction, with CtIP’s DIP-box subsequently replacing BRCA1’s DIP-box, allowing for end resection via MRE11-RAD50. USP1, with its partner UAF1, may then bind to FANCD2 via the UAF1-FANCI interface (*25*), with USP1’s DIP-box subsequently replacing CtIP’s, to remove the ubiquitin signal from FANCD2. However, BRCA1 is also reported to be downstream of FANCD2 in fork protection (*53*) and so there may be multiple mechanisms at play depending on the cellular and environmental context. Regardless, the requirement of the acidic region on FANCD2 to be unoccupied by other interactors for USP1 to efficiently remove the ubiquitin signal may be a protective measure to ensure mono-ubiquitinated FANCD2 has completed its role before the signal is removed.

Different states of FANCD2 may favour recruitment of different interactors. FANCD2 has been reported to have functions independent of FANCI (*51, 54, 55*), and both FANCD2 and FANCI can be mono-ubiquitinated and phosphorylated, resulting in a number of possible states that may exist in the cell. We have shown that CtIP can bind to FANCD2 alone (Figure 2B), mono-ubiquitinated FANCD2 alone (Figure 2A), or mono-ubiquitinated FANCD2 with FANCI (Figure 1D). We also did not observe any global change in the structure of the FANCI-FANCD2 complex, compared to structures in which FAN1 or CtIP is not bound. Furthermore, it has been previously reported that USP1 can bind non-ubiquitinated FANCD2 in pull-down assays (*26*). Overall, these observations suggest that the conserved acidic region is primarily an adaptor site to which other proteins bind as opposed to an allosteric site.

Finally, FANCD2 is implicated in several types of cancer. In lymph node metastases, FANCD2 expression is correlated with poorer prognosis (*56*) and in gliomas, FANCD2 expression is correlated with tumour grade (*57*). Further, FANCD2 has been identified in CRISPR/Cas9 screens to be a potential synthetic lethal target for BRCA1/2 mutant tumours (*58*). The pocket to which the leucine anchor inserts may be a potential target for designing small molecule or peptide inhibitors that block the function of FANCD2.

## Limitations

Protein fragmentation for AlphaFold predictions has been shown to improve sensitivity but it can also reduce specificity (*32*). As such we may have introduced more false positives using this approach, however given the number of sequences that matched our motif criteria but failed to yield relatively high ipTM and PAE scores we believe false positives should be minimal. On the other hand, during our filtering process for FANCD2 interactors, we may have eliminated true FANCD2 interactors. This was to improve computational tractability. We may have also missed supplementary interaction sites outside of the fragment used. The exact fragment used can also affect the ipTM and PAE scores, which we used to cluster into confident and unconfident predictions. We have assumed the variation due to specific fragments lengths is negligible. Further, AlphaFold models are predictions and require experimental validation. We have provided a simple experimental validation for a subset of the candidate interactors. For tractability, we have used peptides encompassing the DIP-box of interactors, guided by AlphaFold predictions. This allowed us to readily use them in our binding and deubiquitination assays, bypassing the need for heterologous production of these large proteins that often exist in even larger assemblies. However, in the context of the full-length interactors in protein complexes it is possible further interactions may be present.

### Conclusions

We have identified a conserved acidic region on FANCD2 that serves as a binding platform for multiple proteins involved in DDR pathways. Through structural and biochemical approaches, we demonstrate that this site mediates direct physical interactions with FAN1, CtIP, BRCA1, and others via a SLiM which we refer to as a DIP-box. The mutually exclusive nature of these interactions, combined with the diversity of predicted interactors spanning end resection, chromatin remodelling, and RNA processing, positions FANCD2 as a hub coordinating multiple aspects of genome maintenance. This adds to the already substantial similarities between FANCD2 and PCNA. Overall, it seems FANCD2’s critical role in ICL repair may be the coordination of multiple sub-pathways needed to manipulate the DNA, through DIP-box interactions.

## Materials and methods

### Protein production

FANCD2 mutations were introduced with site-directed mutagenesis using the Q5 protocol (New England Biolabs). Proteins were expressed as described previously (*17, 25, 27*). Protein purification buffers and columns used are listed in Table S3. Briefly, His_6_-3C-FANCD2 and mutants, His_6_-TEV-USP1^G670A,G671A^, His_6_-3C-UAF1, and His_6_-TEV-V5-FANCI were expressed separately in *Sf*21 insect cells. His_6_-TEV-FAN1^1-71^ (codon optimized for *E. coli*) was expressed in BL21 cells, grown to an OD_600_>0.8 in 2YT media supplemented with 500 µM zinc chloride, via induction with 0.2 mM IPTG and subsequent incubation overnight at 16 °C. Insect and bacterial cells were lysed by sonication, clarified, and proteins purified by Ni-NTA affinity chromatography then, for all proteins except His_6_-TEV-FAN1^1-71^, anion exchange chromatography. At this stage protein aliquots were occasionally flash frozen in liquid nitrogen and stored -80°C. For His_6_-TEV-USP1^G670A,G671A^, TEV protease treatment was performed overnight at ∼1:10 protease to target protein with gentle agitation, resulting in cleaved protein with an N-terminal glycine extension. Subtractive Ni-NTA affinity chromatography was subsequently performed. All proteins were concentrated to 2-10 mg/mL and separated by gel filtration. Purified protein was concentrated to 2-15 mg/mL, flash frozen, and stored at -80°C in 10-20 μL single-use aliquots. All steps were performed on ice or at 4°C and completed within 24-36 hours of lysis. FANCD2 was ubiquitinated and purified using an engineered Ube2T and SpyCatcher-SpyTag setup described in detail elsewhere (*37*). For FANCD2, the His_6_-3C tag was removed by 3C protease treatment during preparation of the mono-ubiquitinated version.

### Peptides

Peptides were ordered from Genosphere in TFA salt at >95% purity. Peptides were dissolved to 1-3mM in milliQ water. The fluorescently labelled CtIP was ordered with a Cy5 fluorophore at the C-terminal lysine. Sequences of peptides used are provided in Table S4.

### CryoEM sample preparation

The FANCI-FANCD2^Ub^-dsDNA-FAN1^1-71^ complex was prepared by mixing the individually prepared components to a final concentration of 6.75 μM His_6_-TEV-V5-FANCI, 6.75 μM FANCD2^Ub^, 13.5 μM DNA, 20 μM His_6_-TEV-FAN1^1-71^ in 10 mM Tris pH 8, 140 mM NaCl, 2% glycerol, 2 mM DTT. The FANCI-FANCD2^Ub^-dsDNA-CtIP^173-193^ complex was prepared by mixing the individually prepared components to a final concentration of 5 μM His_6_-TEV-V5-FANCI, 5 μM FANCD2^Ub^, 9 μM DNA, 20 μM CtIP^173-193^ in 20 mM Tris pH 8, 140 mM NaCl, 2% glycerol, 2 mM DTT. 61 base pair double-stranded DNA was used for both samples (TGATCAGAGGTCATTTGAATTCATGGCTTCGAGCTTCATGTAGAGTCGACGGTGCTGGGAT; IDT). The sample was incubated at room temperature for approximately 5 minutes immediately prior to preparing grids. For the FAN1 samples, UltrAuFoil 1.2/1.3 300 mesh grids were glow discharged at ∼30 mA for 120 secs. For the CtIP samples, QuantiFoil R1.2/1.3 300 mesh grids were glow discharged at 35 mA for 60 secs. A 3.0 μL aliquot of sample was applied. Both grids were blotted for 3.0 secs and vitrified in liquid ethane using a Vitrobot Mark IV (ThermoFisher) operating at ∼95% humidity at 15°C.

### Cryo-EM data collection and processing

Grid screening and data collection was performed at the Scottish Centre for Macromolecular Imaging (SCMI). Data collection was performed on a CryoARM 300 kV microscope (JEOL) equipped with an APOLLO detector (Direct Electron) and Omega Energy Filter (JEOL). For the FAN1 and CtIP datasets, a total of 14,261 and 21,274 movies, respectively, were collected using Beam-Image shift at 60k magnification. Movies were collected at a super-resolution pixel size of 0.52 Å using SerialEM (*59*). An energy filter slit width of 20eV was used. Movies were collected with a total dose of ∼47 e^-^/Å^2^ over 40 frames (FAN1) or ∼60 e^-^/Å^2^ over 50 frames (CtIP).

Subsequent processing was performed in cryoSPARC v4.7 (*60*) (Figure S2 and S3). Patch motion correction with 2x binning, patch CTF estimation, and manual curation was performed. Blob picking, template picking and Topaz (*61*) were used to identify potential particles which were extracted at 300×300 pixels Fourier cropped to 100×100.

For FAN1 dataset, multiple rounds of 2D classification were used for initial cleaning. Ab-initio reconstruction was performed to generate 7 classes. The particles from the class most clearly resembling the FANCI-FANCD2 clamp were then used to generate 3 further classes by ab-initio. Multiple rounds of heterogeneous refinement using these 3 classes were performed to further clean up the particles. Particles corresponding to the FANCI-FANCD2_Ub_-FAN1 volume were re-extracted at 300×300 pixels without Fourier cropping. Additional heterogenous refinement cleaning was performed using the FANCI-FANCD2_Ub_-FAN1 starting model low-pass filtered at 12 Å and one copy at 30 Å. Non-uniform refinement (*62*) yielded a structure with a global resolution of ∼3.5 Å from 264,696 particles. Local refinement was initially performed with a mask covering the FAN1 fragment and the residues of FANCD2 interacting with FAN1. To further improve FAN1 density, 3D classification was performed with 20 classes and a mask only covering the FAN1 fragment. The 20 different classes were examined in ChimeraX and only the classes fully or partially covering the predicted location of the FAN1 fragment (12 out of 20) were selected for a final round of local refinement using a gaussian priors of 2° over rotation and 1 Å over shifts with marginalization and non-uniform refinement.

For the CtIP dataset, multiple rounds of 2D classifications and heterogenous refinement were used in parallel for initial cleaning with selected classes pool and duplicates removed. Further particle cleaning was performed using heterogenous refinement with one starting model corresponding to the FANCI-FANCD2_Ub_-CtIP volume and multiple “junk” starting models distinct from the protein complex of interest, all low pass filtered to 20 Å. Additional heterogenous refinement cleaning was performed using the same starting model low-pass filtered at 12 Å, and two copies at 30 Å. During this cleaning particles were re-extracted at 300×300 pixels without Fourier cropping. Final cleaning steps involved separating particles by 3D Alignment error and 3D Class Probability. Non-uniform refinement (*62*) yielded a structure with a global resolution of ∼4.1 Å from 218,344 particles. Local refinement was performed with a mask covering the helical domain of FANCD2 using a gaussian priors of 2° over rotation and 1 Å over shifts with marginalization and non-uniform refinement.

Model refinement was performed using ISOLDE (*63*), COOT (*64*), and phenix (*65*) using the AlphaFold3 models of FANCD2 with FAN1 and CtIP as starting points. The models were docked in ChimeraX (*66*) using fitmap and the regions with no clear density were pruned. For the FAN1 structure, COOT was used to refine the model into the locally refined sharpened map. For the CtIP structure, ISOLDE and COOT were used to refine the model into the locally refined sharpened map. Finally, phenix.real_space_refine was used to further refine the models and the B-factors against the unsharpened map (FAN1) or sharpened map (CtIP). For FAN1 both Ramachandran and secondary structure restraints were incorporated during phenix refinement. For CtIP the FANCD2-CtIP AlphaFold model was used as a reference model and Ramachandran restraints were incorporated during phenix refinement.

### Fluorescence measurements

Thawed FANCD2 or FANCD2 mutant were exchanged into 2x Assay buffer (40 mM Tris pH 8, 300 mM NaCl, 5% glycerol, 1mM DTT) using Biospin-30 columns (Biorad). Concentration was re-measured using a Nanodrop. CtIP-Cy5 peptide was diluted to 20 μM in MilliQ water, then diluted to 200 nM in 0.05% Tween20. CtIP-Cy5 peptide and a 2-fold serial dilution series of protein were added 1:1 to yield a CtIP-Cy5 concentration of 100 nM and final assay buffer of 20 mM Tris pH 8, 150 mM NaCl, 2.5% glycerol, 0.5 mM DTT, and 0.025% Tween20. Prior to fluorescence measurement samples were briefly centrifuged then transferred into premium capillaries (NanoTemper Technologies). Measurements were performed at 22 °C on a Monolith NT.115 instrument (NanoTemper Technologies) using the red channel. Laser power was set to 10%. Buffer exchange and titrations were performed as technical duplicates for each mutant.

#### Model fitting was performed using a one-binding site model

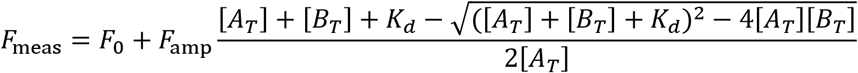

Where *F*_meas_ is the fluorescence measured, *F*_0_ is baseline, *F*_amp_ is the maximum amplitude of the fluorescence change, [*A*_*T*_] is the constant concentration of the fluorescent binding molecule (CtIP peptide), and [*B*_*T*_] is the varying concentration of the other binding molecule (FANCD2). Given that the mutants did not go to saturation, it was assumed that *F*_amp_ was the same for each titration. The fitting was performed to the combined datasets using Markov Chain Monte Carlo analysis (as implemented in https://github.com/mlrennie/pytrate) and assuming a uniform uncertainty of 20 fluorescence units in the experimental measurements. The distributions and correlations between of all the fitted parameters are shown in Figure S8.

### Deubiquitination assays

Deubiquitination assays were performed as previously described (*17, 25*). A 2x substrate mix and a 2x enzyme mix were prepared separately and mixed 1:1 to initiate the reaction. Both mixes were setup on ice and then incubated at room temperature for at least 20 min prior to reaction initiation and during the reaction.

For competition assays, the 2x substrate mix was prepared by diluting stocks of FANCD2^Ub^ with DUB assay buffer (20 mM Tris pH 8.0, 75 mM NaCl, 5% glycerol, 1 mM DTT) to yield a concentration of 2 μM FANCD2^Ub^ substrate and 8 μM peptide or His_6_-TEV-FAN1^1-71^. The 2x enzyme mixes were prepared by diluting stocks of USP1, His_6_-3C-UAF1, and peptide with DUB buffer to yield a concentration of 200 nM USP1-UAF1 enzyme.

For deubiquitination assays with FANCD2 mutants a DUB-step procedure was used to generate the 2x substrate mix (*17, 26*). This involved first mono-ubiquitinating wild-type control or mutant FANCD2 with the ubiquitination machinery including an enhanced Ube2T (*38*). Reactions were setup with 4 µM His_6_-3C-FANCD2, 100 nM Uba1, 4.4 μM Ube2Tv4, 4.4 μM FANCL^109-375^, 8 µM ubiquitin, 5 mM MgCl_2_, 2.5 mM ATP, prepared by diluting all components into DUB assay buffer. Reactions were incubated for 45min at room temperature before being terminated with ∼7 U/mL apyrase (New England Biolabs) and subsequent incubation at room temperature for 5 min. The reaction mix was then diluted 2-fold with DUB assay buffer to give a 2x substrate mix. The 2x enzyme mixes were prepared by diluting stocks of USP1 and His_6_-3C-UAF1 with DUB buffer to yield a concentration of 200 nM USP1-UAF1 enzyme.

Aliquots of 4 μL of reaction were terminated at the indicated timepoints by addition of 20 μL 1.2x NuPAGE LDS buffer (Thermo Fisher) supplemented with 120 mM DTT. SDS-PAGE was performed using Novex 4–12% Tris-glycine gels and subsequent staining of the gels with Instant-Blue Coomassie stain (Expedeon). Gels were imaged on an Odyssey CLx (LI-COR) using the 800-nm channel.

### Cell survival assays

U2OS cells and U2OS FANCD2 knockout cells were purchased from Ubigene Biosciences and were grown in Dulbecco’s Modified Eagle Medium (DMEM) supplemented with 10% fetal calf serum, 100 U/ml penicillin and 100 µg/ml streptomycin. Wild-type or mutant FANCD2-EGFP plasmids were transfected by Lipofectamine 3000 Transfection Reagent (Thermofisher) according to manufacturer’s instructions. Transfected cells were treated with MMC for 16 hours, washed with PBS twice, and incubated on six-well plates for 7 days. Cells were then fixed by 4% paraformaldehyde and stained by 0.1% crystal violet. Staining was quantified using an Odyssey CLx (LI-COR). Each treatment was normalised relative to the 0 ng/mL of that treatment. Western blots were performed using anti-FANCD2 (MRC PPU Reagents and services 33539) and anti-tubulin (Proteintech 66031-1-lg) (Figure S9). Mitomycin C was purchased from Roche (10107409001).

### Motif searching for FANCD2 interactors

We hypothesized that key features of the interaction include:

a. The anchor Leu
b. An Arg or Ser should follow the Leu
c. The Leu should not be preceded by a Pro
d. At least 4 Arg or Lys residues within a 7 amino acid window either side of the Leu
e. At least 2 Arg or Lys residues in a row within the 7 amino acid window
f. A maximum of 2 Asp or Glu within the 7 amino acid window
g. No Asp or Glu within the 3 amino acids either side of the Leu
h. No aromatic residues within the 3 amino acids either side of the Leu
i. An Arg within the 3 amino acids before the leucine
j. A maximum of 2 hydrophobic residues within the 3 amino acids either side of the Leu

From UniProt, the reviewed homo sapiens protein sequences with any evidence for nucleus subcellular location were downloaded on 22^nd^ May 2025. Of these 5,700 sequences, 479 had at least one region matching the criteria above. For these hits, AlphaFold models from the AFDB were used to predict if the region had a pLDDT score less than 70, which correlates with disordered regions. A total of 243 sequences from 210 proteins met these criteria. To narrow this further, the UniProt function text for each of the hits was searched for terms matching “DNA repair”, homologous recombination”, “double-strand break”, “DSB”, or “DNA damage”. The resulting 47 sequences across 39 proteins were extracted with approximately 50 residues either side of the motif and passed through the AlphaFold3 server (*31*) with full-length FANCD2, using the default settings. This resulted in 40 predictions due to overlap of some the of sequences within the same protein. Python scripts to perform much of this processing were developed with the assistance of Claude Sonnet 4 and Opus 4. Models were visualized using ChimeraX (*66*).

### Statistical analysis

To generate a threshold for the ipTM and minPAE score respectively, we used K-means clustering with k=2 to estimate the means of confident prediction and non-confident prediction clusters. We took the average of these means as the threshold for separating the clusters and only if a prediction passed both the ipTM and minPAE thresholds was it considered a confident prediction. The ipTM and minPAE thresholds were 0.33 and 12.40 Å, respectively.

### Conservation

A sequence alignment for FANCD2 was obtained from the ProViz server (Quest for Orthologues alignment) (*41*). FANCD2 was colored by the AL2CO (*40*) entropy-based measure as implemented in ChimeraX.

### Electrostatic calculations

PDB2PQR and APBS were used for electrostatic calculations via the webserver (https://server.poissonboltzmann.org/) (*42*). Default settings were used aside from the ions for which +1 ions with radius 2.0 and -1 ions with radius 1.8 were included, both at concentrations of 0.15 M. Electrostatics were visualized in ChimeraX (*66*).

## Acknowledgements

We thank past and current members of the Walden laboratory for experimental suggestions, comments on the manuscript and their support. The authors acknowledge the Neil Bulleid Integrated Protein Analysis facility for access to MST. We acknowledge the Scottish Centre for Macromolecular Imaging (SCMI) for assistance with cryo-EM experiments and access to instrumentation, funded by the MRC (MC_PC_17135, MC_UU_00034/7, MR/X011879/1) and SFC (H17007). The authors thank Mark Meenan, Paul McLaughlin, and Iain Sim for maintenance of the GPU server running cryoSPARC. H.W., M.L.R., and Z.C. were supported by a Medical Research Council grant to H.W. (MR/W025256/1).

## Author contributions (CRediT)

Conceptualization – M.L.R., H.W.; Investigation – M.L.R., Z.C., G.B., J.S., C.I.; Supervision – M.L.R., H.W.; Data Curation – M.L.R., Z.C.; Visualization – M.L.R., Z.C.; Writing (original draft) – M.L.R.; Writing (review and editing) – M.L.R., Z.C., C.I., H.W.

## Data Availability

The atomic coordinates have been deposited to the PDB under accession codes 28UE (CtIP) and 28UF (FAN1). The cryo-EM maps have been deposited to the EMDB under accession codes EMD-56824 (CtIP) and EMD-56825 (FAN1).

