## Supplementary Figures and Tables for "Discovery of a FANCD2-interacting protein motif (DIP-box) linking DNA Damage Response processes"

**Title:**

Affiliations:

Affiliations:

Table of Contents

**SUPPLEMENTARY FIGURES AND TABLES**

**Figure S1**........ **S2**
**Figure S2**........ **S3**
**Figure S3**........ **S4**
**Figure S4**........ **S5**
**Figure S5**........ **S6**
**Figure S6**........ **S7**
**Figure S7**........ **S9**
**Figure S8**........ **S10**
**Figure S9**........ **S11**
**Table S1**........ **S12**
**Table S2**........ **S14**
**Table S3**........ **S17**
**Table S4**........ **S18**

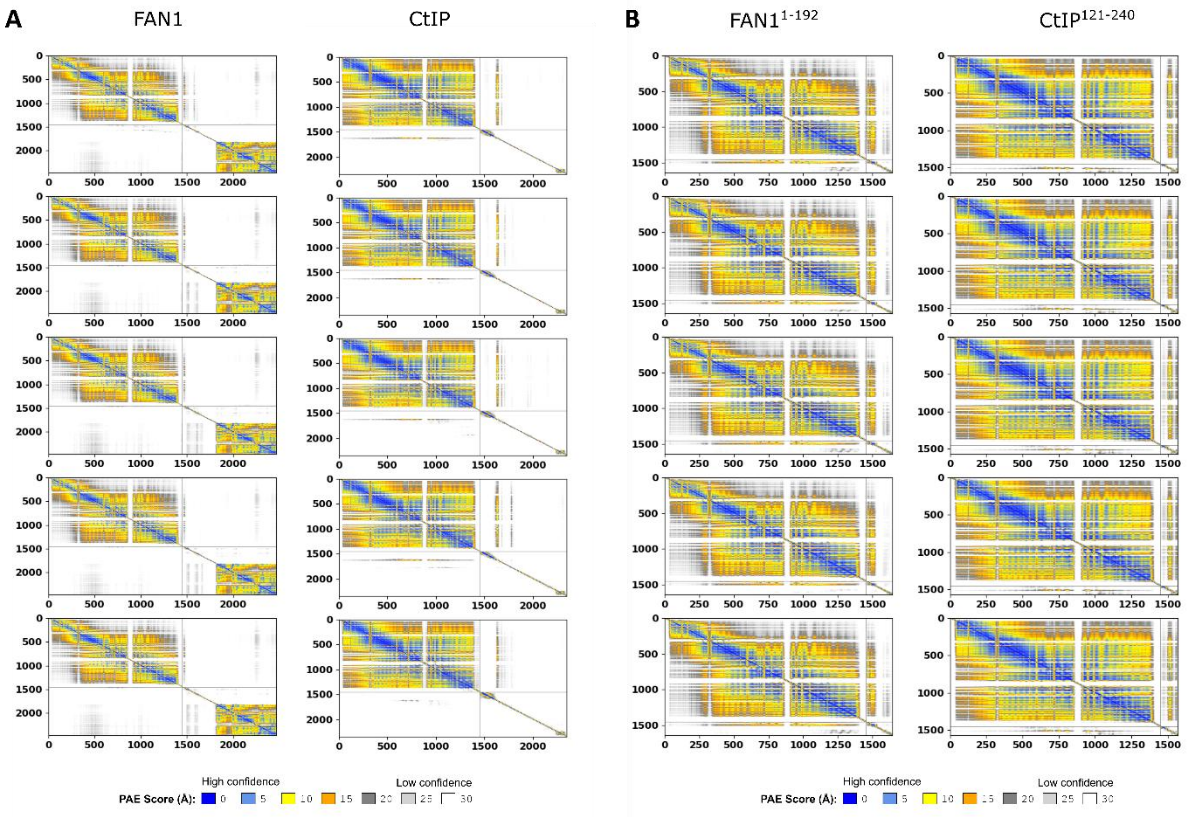


**Figure S1:** PAE plots for FANCD2 with FAN1 and with CtIP. (A) Full-length CtIP and FAN1. (B) Fragments of FAN1 and CtIP. The five replicates generated by AlphaFold3 are shown. Default settings were used.

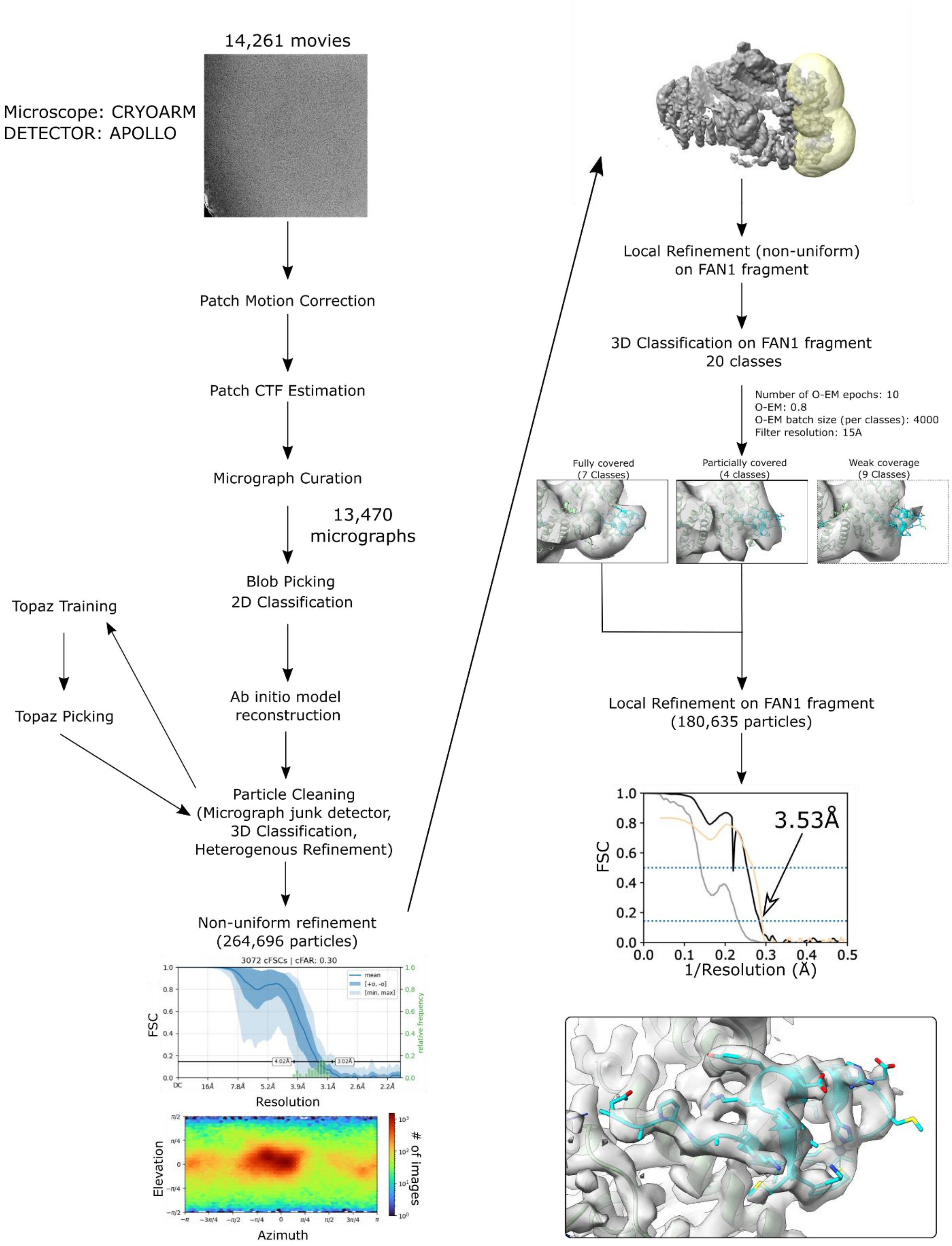


**Figure S2:** Cryo-EM particle processing workflow for the FAN1-bound structure. FSC and particle distributions for the consensus refinement are shown on the bottom left. Half-map FSCs for no mask (dark gray) and tight mask with correction by noise substitution (black) are shown for the final reconstruction. Model-map FSC is shown in tan. The refined structure, fit within the map, is shown for the UBZ binding site.

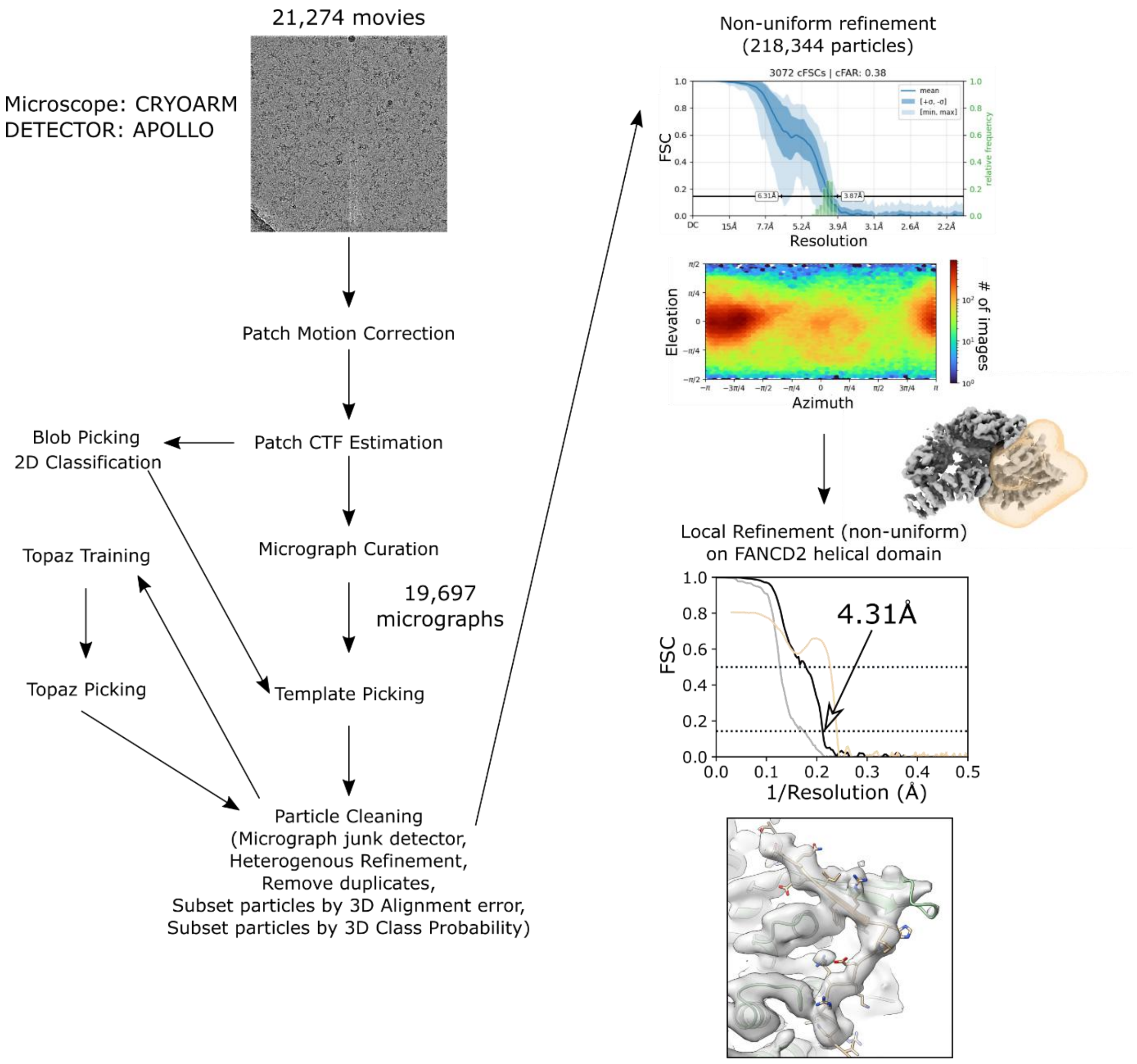


**Figure S3:** Cryo-EM particle processing workflow for the CtIP-bound structure. FSC and particle distributions for the consensus refinement are shown on the top right. Half-map FSCs for no mask (dark gray) and tight mask with correction by noise substitution (black) are shown for the final reconstruction. Model-map FSC is shown in tan. The refined structure, fit within the map, is shown for the CtIP binding site.

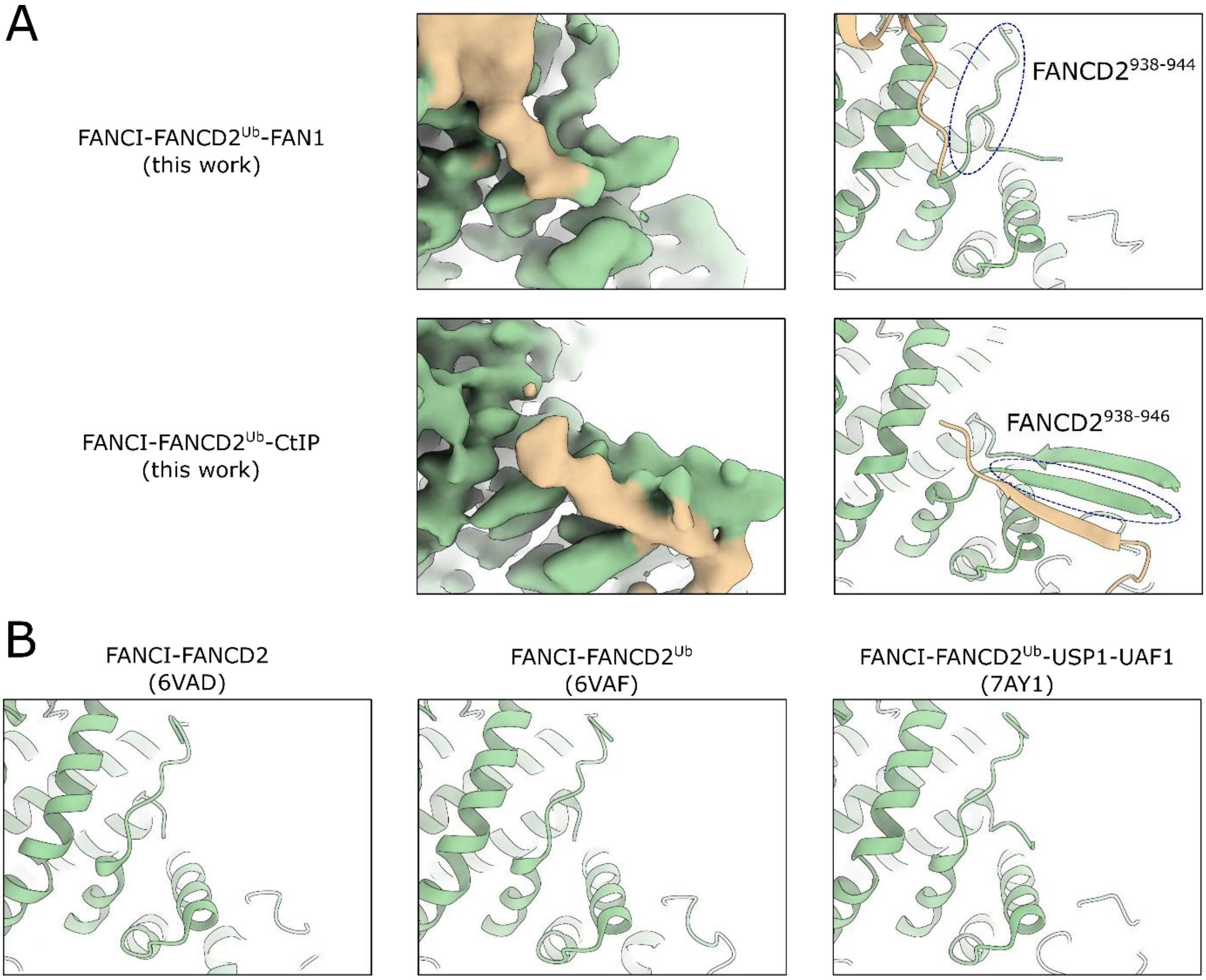


**Figure S4:** Comparison of experimental structures of FANCD2 in different forms. Only in the presence of CtIP, FANCD2 residues 938-944 restructure and residues 945-958 become ordered and together form an antiparallel β-sheet. Residues undergoing conformational change are highlighted with dashed ovals.

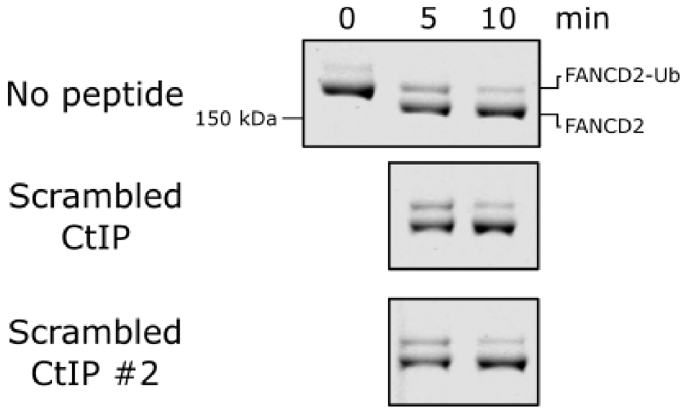


**Figure S5:** Competition deubiquitination assays with 50 µM scrambled CtIP peptides.

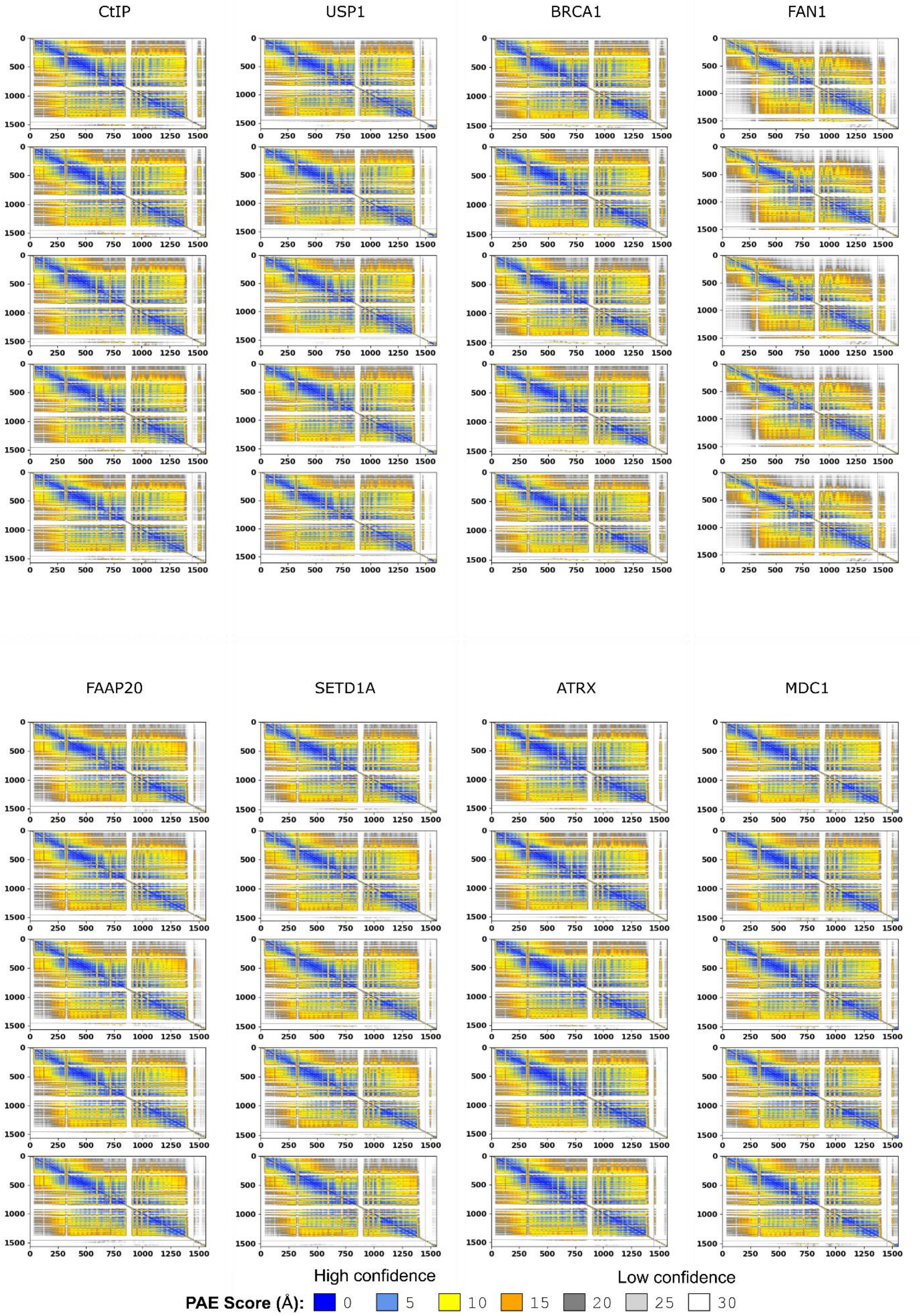
CtIP
USP1
BRCA1
FAN1
FAAP20
SETD1A
ATRX
MDC1
High confidence
Low confidence
PAE Score (Å): 0 5 10 15 20 25 30

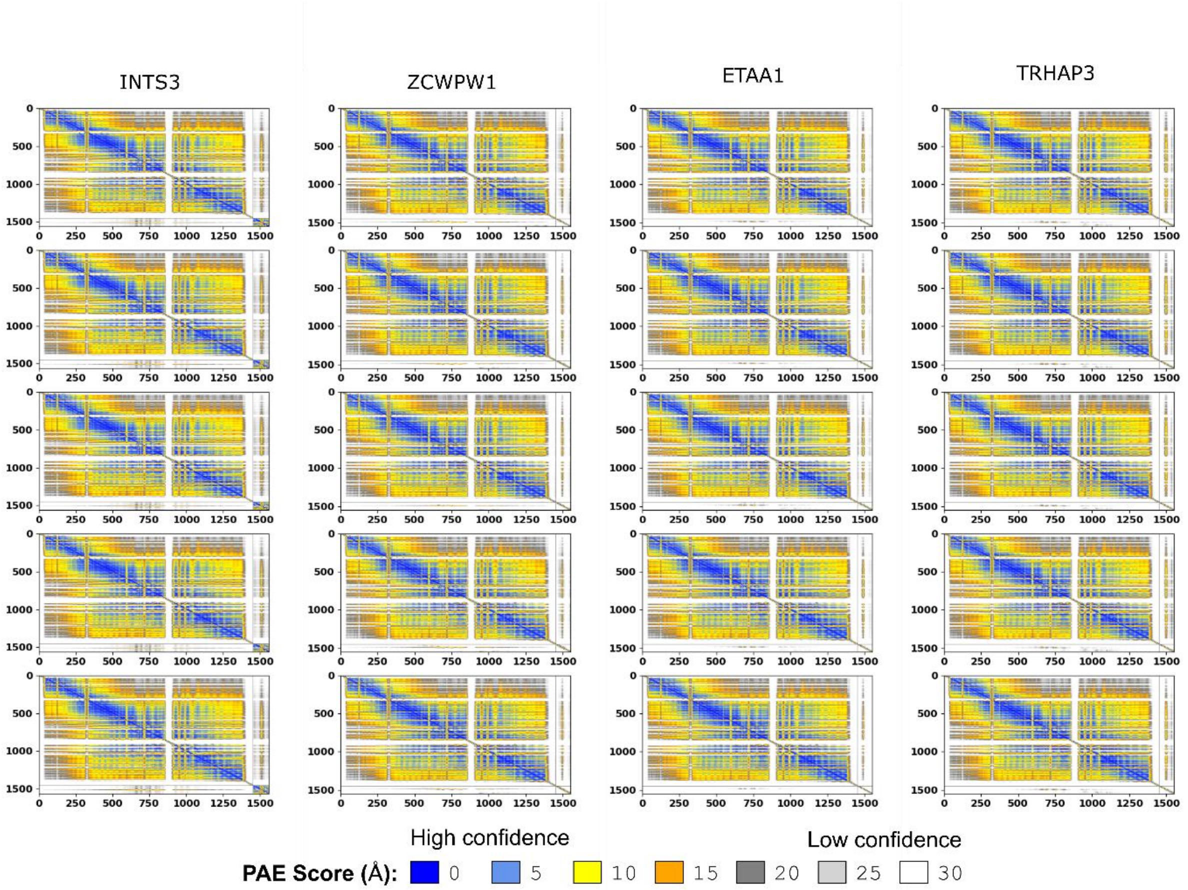


**Figure S6:** PAE plots for candidate FANCD2 interactors. Interactors are indices 1451 onwards. The five replicates generated from AlphaFold3 are shown. CtIP and FAN1 fragments are from Figure S1.

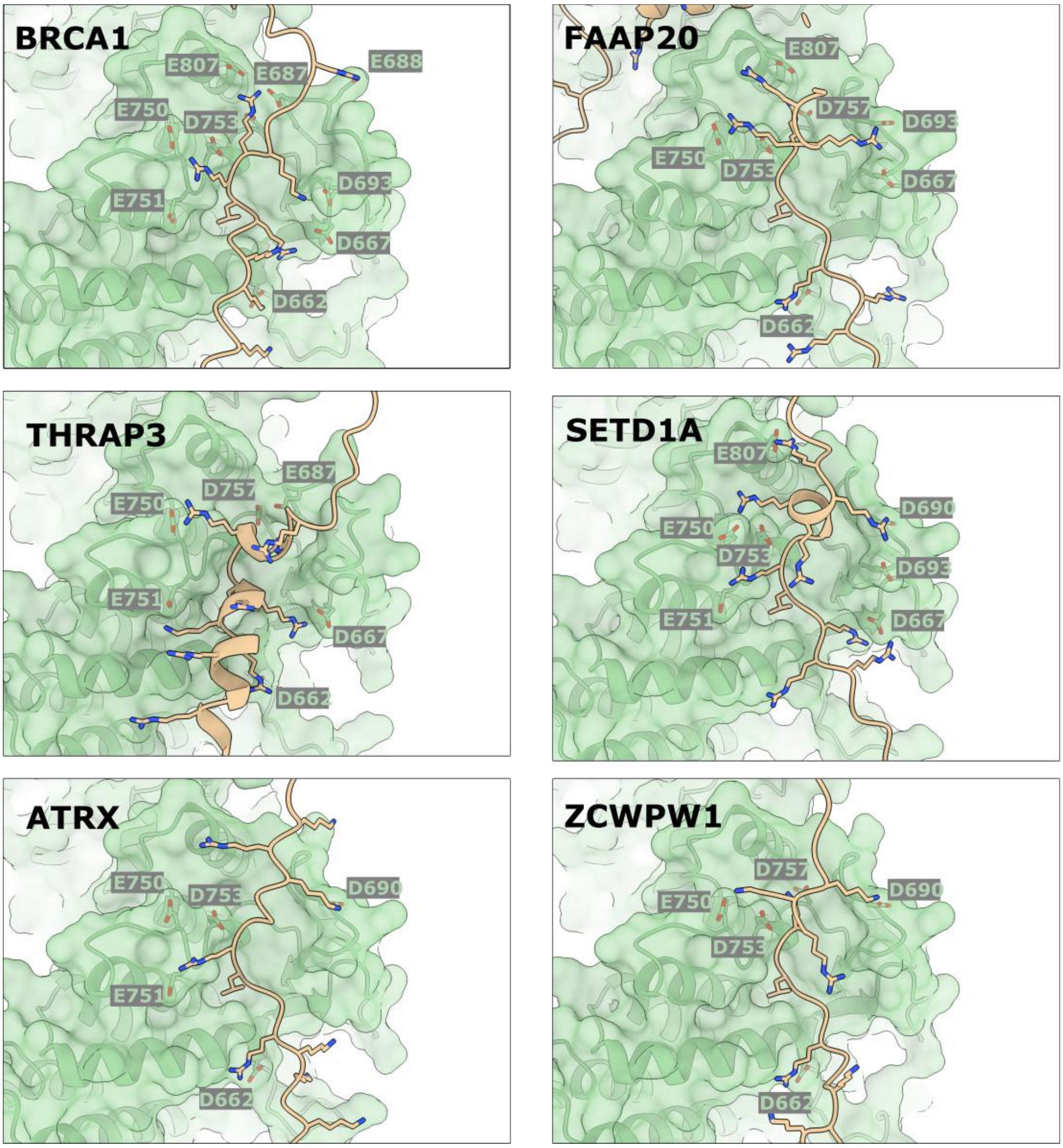


**Figure S7:** Predicted atomic models of candidate DIP-boxes. Acidic residues of FANCD2 (green) are labelled. Arginines and lysines of each interactor, as well as the leucine anchor are shown.

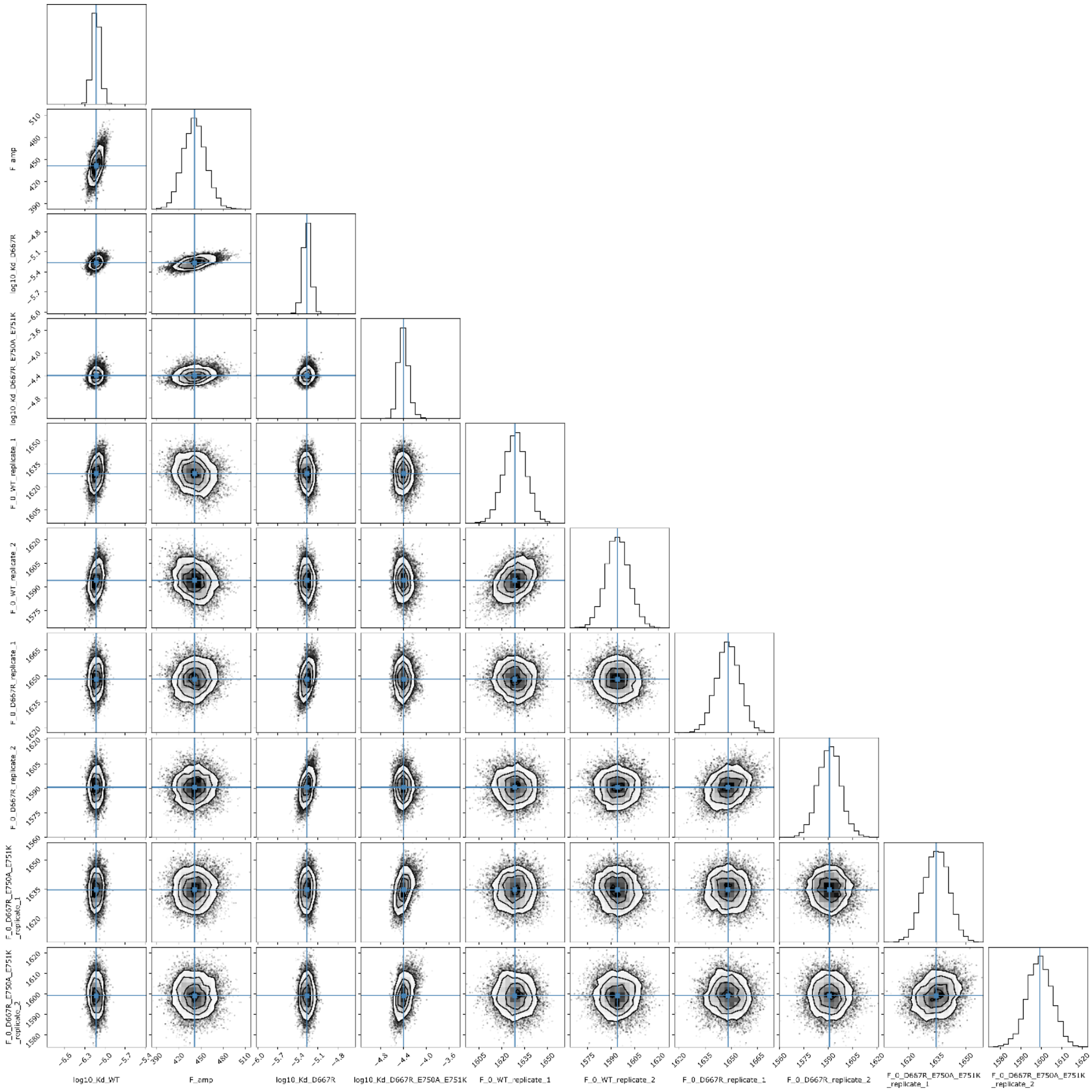

**Figure S8:** Corner plot showing parameter correlations for fitting to the fluorescence data in Figure 1D-E.

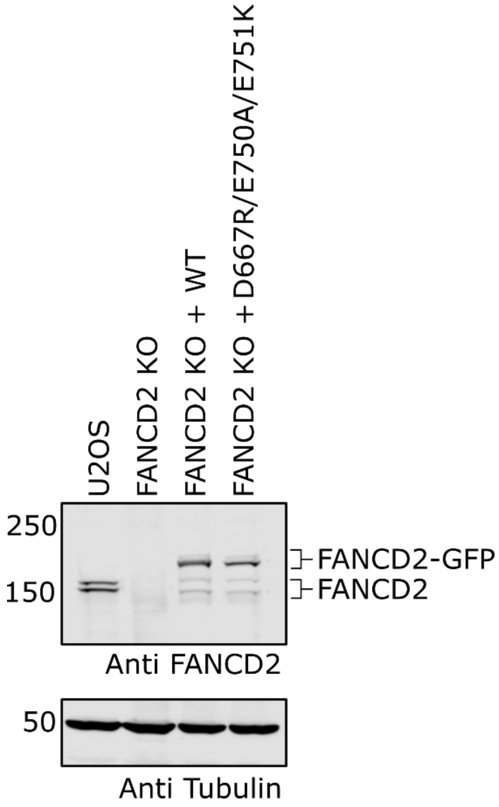


**Figure S9:** Western blots of whole cells after transfection but prior to MMC treatment.

**Table S1.** Cryo-EM data collection and model refinement statistics

| | FAN1 Consensus | FAN1 Focused | CtIP Consensus | CtIP Focused |
|---|---|---|---|---|
| **Data collection and processing** | | | | |
| Microscope | CryoARM (JEOL) | | | |
| Detector | APOLLO (Direct Electron) | | | |
| Nominal Magnification | 60,000 | | | |
| Voltage (kV) | 300 | | | |
| Electron Dose ($e^-/Å^2$) | ~47 | | ~60 | |
| Defocus range (µm) | ~0.9-2.5 | | ~0.5-2.0 | |
| Pixel Size (Å) | 0.52 | | 0.516 | |
| Symmetry imposed | C1 | | C1 | |
| Map resolution (Å) | 3.50 | 3.53 | 4.11 | 4.31 |
| FSC threshold | 0.143 | 0.143 | 0.143 | 0.143 |
| Map resolution range (Å)[a] | - | 3.1-11 | - | 3.7-14 |
| FSC threshold | - | 0.143 | - | 0.143 |
| EMDB ID | EMD-56825 (additional map) | EMD-56825 | EMD-56824 (additional map) | EMD-56824 |
| **Refinement** | | | | |
| Initial models used | - | AlphaFold | - | AlphaFold |
| Map sharpening B-factor ($Å^2$) | - | 124.9 | - | 175.8 |
| Q-score | - | 0.464 | - | 0.345 |
| Bond length rmsd (Å) | - | 0.003 | - | 0.002 |
| Bond angle rmsd (°) | - | 0.436 | - | 0.443 |
| All-atom clashscore | - | 5.65 | - | 4.58 |
| Ramachandran plot | - | | | |
| Outliers (%) | - | 0.00 | - | 0.00 |

| | | | | |
|---|---|---|---|---|
| Allowed (%) | - | 2.44 | - | 3.30 |
| Favored (%) | - | 97.56 | - | 96.70 |
| Rama-Z (whole) | - | 1.61 | - | 2.05 |
| CaBLAM Outliers (%) | - | 0.79 | - | 0.92 |
| Rotamer outliers (%) | - | 0.39 | - | 0.32 |
| PDB ID | - | 28UF | - | 28UE |

[a]1% and 99% quantiles from local resolution computed in cryoSPARC with Adaptive Window Factor of 10

**Table S2.** Fragments used for AlphaFold3 predictions with FANCD2.

| Uniprot ID | Gene Name | Predicted Leu anchor | Fragment used | ipTM score | minPAE score |
|---|---|---|---|---|---|
| O15047 | SETD1A | 1394 | 1351-1450 | 0.39 | 6.12 |
| O15297 | PPM1D | 45 | 1-100 | 0.22 | 19.45 |
| O15297 | PPM1D | 555 | 491-C-terminus | *** | *** |
| O15297 | PPM1D | 584 | 491-C-terminus | 0.33 | 7.86 |
| O75417 | POLQ | 2171 | 2131-2230 | 0.3 | 9.79 |
| O94761 | RECQL4 | 369 | 311-430 | 0.23 | 18.03 |
| O94761 | RECQL4 | 374 | 311-430 | *** | *** |
| O94782 | USP1 | 23 | 1-150 | 0.41 | 4.24 |
| O95071 | UBR5 | 288 | 231-340 | 0.2 | 20.54 |
| P00519 | ABL1 | 1012 | 961-1060 | 0.24 | 15.48 |
| P28715 | ERCC5 | 1176 | 1130-C-terminus | 0.32 | 8.99 |
| P38398 | BRCA1 | 611 | 561-660 | 0.47 | 2.22 |
| P46100 | ATRX* | 1140 | 1091-1200 | 0.39 | 7.81 |
| P46100 | ATRX | 1189 | 1131-1250 | 0.48 | 3.9 |
| Q14676 | MDC1 | 1881 | 1831-1930 | 0.41 | 4.42 |
| Q14839 | CHD4 | 346 | 291-400 | 0.22 | 18.25 |
| Q56NI9 | ESCO2 | 7** | 1-100 | 0.22 | 19.69 |

| | | | | | |
|---|---|---|---|---|---|
| Q68E01 | INTS3 | 909 | 851-960 | 0.37 | 5.9 |
| Q6IN85 | PPP4R3A | 663 | 601-720 | 0.27 | 18.41 |
| Q6NZ36 | FAAP20 | 11 | 1-110 | 0.42 | 5.96 |
| Q70CQ1 | USP49 | 117 | 61-170 | 0.23 | 16.53 |
| Q86YC2 | PALB2 | 171 | 121-220 | 0.27 | 12.53 |
| Q8N9B5 | JMY | 875 | 821-930 | 0.2 | 22.01 |
| Q8NFZ0 | FBH1 | 152 | 101-200 | 0.19 | 20.41 |
| Q96NY9 | MUS81 | 7 | 1-100 | 0.27 | 18.98 |
| Q96QE3 | ATAD5 | 520 | 471-570 | 0.31 | 11.85 |
| Q99708 | RBBP8 (CtIP) | 176 | 121-240 | 0.49 | 2.83 |
| Q99728 | BARD1 | 328 | 271-380 | 0.19 | 21.36 |
| Q9BUA3 | SPINDOC | 196 | 131-250 | 0.23 | 16.8 |
| Q9BZF9 | UACA | 4** | 1-100 | 0.21 | 25.25 |
| Q9BZF9 | UACA | 8 | 1-100 | *** | *** |
| Q9H0M4 | ZCWPW1 | 203 | 151-250 | 0.41 | 5.67 |
| Q9H6W3 | RIOX1 | 43 | 1-100 | 0.27 | 14.23 |
| Q9H9Q4 | NHEJ1 | 286 | 201-C-terminus | 0.21 | 16.71 |
| Q9NP87 | POLM | 2** | 1-100 | 0.26 | 22.3 |
| Q9NS56 | TOPORS | 35 | 1-100 | 0.3 | 6.72 |

| | | | | | |
|---|---|---|---|---|---|
| Q9NS56 | TOPORS | 725 | 671-780 | 0.21 | 20.69 |
| Q9NY74 | ETAA1 | 37 | 1-100 | 0.34 | 5.82 |
| Q9NZ56 | FMN2 | 84 | 31-140 | 0.27 | 14.31 |
| Q9UIG0 | BAZ1B | 1282 | 1231-1330 | 0.19 | 17.58 |
| Q9UMN6 | KMT2B | 77 | 21-130 | 0.2 | 19.61 |
| Q9Y2K7 | KDM2A | 14 | 1-100 | 0.25 | 20.7 |
| Q9Y2K7 | KDM2A | 385 | 331-440 | 0.29 | 16.31 |
| Q9Y2M0 | FAN1 | 17 | 1-192 | 0.5 | 3.61 |
| Q9Y2S7 | POLDIP2 | 51 | 1-100 | 0.21 | 24.06 |
| Q9Y2W1 | THRAP3 | 37 | 1-100 | 0.41 | 4.28 |
| Q9Y2W1 | THRAP3 | 45 | 1-100 | *** | *** |

*model has L1189 within hydrophobic pocket
**close to the N-terminus and so does not satisfy the criteria of the N-terminal side of the leucine
***Leucine anchor was very close to another hit sequence and so single fragment was used

**Table S3.** Protein purification buffers.

| | Purification step (column)/Experiment | Buffer composition |
|---|---|---|
| USP1 or UAF1 | Lysis | 50 mM Tris pH 8.0, 150 mM NaCl, 5% (v/v) glycerol, 5 mM β-mercaptoethanol, 10 mM Imidazole, 2 mM $MgCl_2$, 1x cOmplete EDTA-free protease inhibitor cocktail, >10 units/mL benzonase |
| | Ni-NTA Wash 1/Subtractive | 50 mM Tris pH 8.0, 500 mM NaCl, 5% (v/v) glycerol, 5 mM β-mercaptoethanol, 10 mM Imidazole |
| | Ni-NTA Wash 2 | 50 mM Tris pH 8.0, 100 mM NaCl, 5% (v/v) glycerol, 1 mM TCEP, 10 mM Imidazole |
| | Ni-NTA Elution | 50 mM Tris pH 8.0, 75 mM NaCl, 5% (v/v) glycerol, 1 mM TCEP, 250 mM Imidazole |
| | Anion Exchange (ResourceQ 1 mL) | 50 mM Tris pH 8.0, 5% (v/v) glycerol, 1 mM TCEP, 100-1000 mM NaCl |
| | Gel Filtration (Superdex 200 Increase 10/300 GL) | 20 mM Tris pH 8.0, 150 mM NaCl, 5% (v/v) glycerol, 5 mM DTT |
| FANCD2, FANCI | Lysis | 50 mM Tris pH 8.0, 400 mM NaCl, 5% (v/v) glycerol, 5 mM β-mercaptoethanol, 10 mM Imidazole, 2 mM $MgCl_2$, 1x cOmplete EDTA-free protease inhibitor cocktail, >10 units/mL benzonase |
| | Ni-NTA Wash 1 | 50 mM Tris pH 8.0, 400 mM NaCl, 5% (v/v) glycerol, 5 mM β-mercaptoethanol, 10 mM Imidazole |
| | Ni-NTA Wash 2 | 50 mM Tris pH 8.0, 150 mM NaCl, 5% (v/v) glycerol, 1 mM TCEP, 10 mM Imidazole |
| | Ni-NTA Elution | 50 mM Tris pH 8.0, 100 mM NaCl, 5% (v/v) glycerol, 1 mM TCEP, 250 mM Imidazole |
| | Anion Exchange (HP Q 5 mL) | 50 mM Tris pH 8.0, 5% (v/v) glycerol, 1 mM TCEP, 100-1000 mM NaCl |
| | Gel Filtration (Superose 6 Increase 10/300 GL) | 20 mM Tris pH 8.0, 400 mM NaCl, 5% (v/v) glycerol, 5 mM DTT |
| FAN1 | Lysis | 1x Phosphate buffered saline (PBS), 2 mM β-mercaptoethanol, 2 mM $MgCl_2$, 1x cOmplete EDTA-free protease inhibitor cocktail, >10 units/mL benzonase |
| | Ni-NTA Wash | 1x PBS, 0.4 mM TCEP |
| | Ni-NTA Elution | 1x PBS, 0.4 mM TCEP, 250 mM Imidazole |
| | Gel Filtration (Superdex 75 Increase 10/300 GL) | 20 mM Tris pH 8.0, 150 mM NaCl, 5% (v/v) glycerol, 0.4 mM DTT |

**Table S4.** Peptide sequences ordered from Genosphere.

| Peptide | Sequence |
|---|---|
| CtIP$^{173-193}$ | VNRLRRKENPHVRYIEQTHTK |
| CtIP$^{173-193}$-Cy5 | VNRLRRKENPHVRYIEQTHT[K-cy5]-[nh2] |
| Scrambled CtIP | VNRERYKRNLHERPIVQTHTK |
| Scrambled CtIP #2 | VNRPRYEKNLHERRIVQTHTK |
| USP1$^{16-33}$ | SPSKKNRLSLKFFQKKET |
| BRCA1$^{606-625}$ | PKKNRLRRKSSTRHIHALEL |
| ATRX$^{1179-1198}$ | VIVKEKKRNSLRTSTKRKQA |
| FAAP20$^{1-20}$ | MEAARRPRLGLSRRRPRPAG |
| THRAP3$^{35-54}$ | RSLSRSRKRRLSSRSRSRSY |
| MDC1$^{1871-1890}$ | EEPNRIPSRSLRRTKLNQES |
| ZCWPW1$^{193-212}$ | PSKKKSNRLTLSKRKKEAHE |
| SETD1A$^{1384-1403}$ | DGEGALRRRSLRSHARRRRP |
| INTS3$^{899-918}$ | NNSLPRKRQSLRSSSSKLAQ |
| ETAA1$^{27-46}$ | GSVVEPGRRRLRSARGSWPC |
| BARD1$^{318-337}$ | NNGKRGHHNRLSSPISKRCR |
